# Sperm whale foraging combines persistent search with flexible prey-capture tactics

**DOI:** 10.64898/2026.07.29.741536

**Authors:** Lorenzo Zaffina, Corrado Monti, Simone Poetto, Maxime Lucas, Henrique João Machado Borges, Raj Deshpande, Pernille Tønnesen, David F. Gruber, Shane Gero, Andrea Santoro, Giovanni Petri

## Abstract

Predators searching patchy environments must coordinate movements across scales, yet this behavioural hierarchy is not yet technically possible to observe in the deep ocean. Here we show that sperm-whale foraging is organized across two nested levels: directionally persistent search paths and flexible prey-capture tactics. We combine acoustic recordings with reconstructed three-dimensional trajectories from 34 sensor-tag deployments on 20 individual whales in both the eastern Caribbean and the mid-Atlantic. Complete foraging paths exhibit heavy-tailed step lengths and superdiffusive displacement, consistent with Lévy-like search. Within these paths, echolocation buzzes resolve into two recurrent acoustic–kinematic tactics. Type A involves brief ramp-ups, slower and more oscillatory movement, and longer post-buzz pauses. Type B involves prolonged ramp-ups, faster and vertically persistent movement, greater roll, slightly straighter approaches, and an energy-model estimate approximately seven times higher. Every sampled whale used both tactics and switched between them within dives, showing that the contrast is not driven by specialization among individuals. An Azorean deployment with prey-field echograms links the tactics to local prey conditions, while independent Dominica deployments recover their core kinematic contrast. Together, these results reveal a foraging hierarchy in which flexible capture actions unfold within persistent, deliberate large-scale search.

## INTRODUCTION

Predators foraging in heterogeneous environments must solve two coupled problems: how to search across space and how to act once prey is encountered. Search determines the sequence of encounters available to an individual, whereas local pursuit and capture determine how it responds to each opportunity. These processes operate over different spatial and temporal scales, but are rarely observed together, particularly in the deep ocean where both predators and prey remain largely inaccessible to direct observation.

Sperm whales (*Physeter macrocephalus*) provide an unusual opportunity to connect these scales. They routinely perform dives exceeding 600 m for more than 40 minutes to exploit mesopelagic and bathypelagic prey [3]. During these dives, whales navigate and hunt via echolocation [4, 5], an ancient odontocete adaptation for which anatomical evidence extends to Oligocene taxa approximately 32 million years ago [6]. Individual prey-capture attempts are marked by *buzzes*: rapid click sequences accompanying the transition from long-range search to close-range target tracking [3, 7, 8]. Multi-sensor biologging tags can therefore align an acoustic marker of prey capture with the movement that precedes, accompanies and follows each attempt.

Buzzes were initially treated primarily as binary indicators of feeding activity [7, 9]. Subsequent studies showed that they are embedded within an information-driven hunting sequence: whales adjust their acoustic gaze across prey layers, select targets from the surrounding prey field and perform rapid manoeuvres during the final approach [10–14]. These observations establish that capture attempts vary acoustically and kinematically. It remains unclear, however, whether this variation resolves into recurrent prey-capture tactics, whether individual whales switch between such tactics within a dive, and how these local actions are embedded within the larger-scale organization of search.

Optimal Foraging Theory (OFT) predicts that generalist predators should vary their behavioural investment as encounter conditions change rather than deploy a single capture strategy [15–17]. Depth-dependent foraging regimes have been described in male sperm whales [10, 18, 19], but much less is known about tactical variation within dives in females and immature animals, which constitute the stable social units of sperm-whale societies. More generally, trajectory-scale search and event-scale prey capture have not been integrated within a common quantitative description of sperm-whale foraging.

Here, we show that sperm-whale foraging is organized across two coupled scales: persistent search over complete trajectories and flexible prey-capture tactics deployed during individual encounters. We do this by analysing synchronized acoustic recordings and reconstructed three-dimensional trajectories from 34 suction-cup sensor tag deployments on 20 predominantly female and immature sperm whales in the Eastern Caribbean [1, 20]. At the trajectory scale, foraging paths exhibit heavy-tailed step lengths and strongly persistent, superdiffusive movement consistent with Lévy-like search. At the scale of individual capture attempts, echolocation buzzes resolve into two recurrent acoustic–kinematic tactics that occur within the same individuals and switch non-randomly within dives; their core behavioural contrast is recovered in an Azorean record with concurrent prey-field echograms [21] and in independent 2024 Dominica deployments.

## RESULTS

Animal movement is organized across nested spatial and temporal scales, with trajectory-scale displacement emerging from sequences of local movements and behavioural decisions [22]. To investigate foraging across trajectory and prey-capture scales, we analysed 34 DTAG deployments collected by the Dominica Sperm Whale Project (DSWP) off the coast of Dominica between 2014 and 2018 [1]. These multi-sensor tags provided continuous depth, high-resolution accelerometry, magnetometry, and audio, enabling three-dimensional reconstruction of underwater movement alongside identification of acoustic-foraging events. Although the acoustic and movement data streams were sampled at different rates, both were logged against a shared internal DTAG clock. Acoustic events could therefore be directly matched to their kinematic context, without additional post-hoc synchronization. In total, the dataset contains 268 hours of recording across 20 distinct individuals, capturing 245 deep foraging dives. For a subset of whales, multiple deployments were available, enabling assessment of within-individual behavioral consistency across deployments and dates. GPS position recorded from the accompanying vessel was available for a subset of deployments (see SI Fig. S1 for an overview of the basic statistics of the DSWP DTAG dataset).

### 3D reconstruction and trajectory-scale organization

To recover the paths within which prey-capture attempts occurred, we reconstructed three-dimensional trajectories using pitch-adaptive dead reckoning based on pressure-derived depth and sensor-derived body orientation at 25 Hz [7, 23] (see Methods). This provided continuous estimates of position, swim speed and orientation for the trajectory- and buzz-scale analyses. Among four reconstruction methods evaluated against vessel-based GPS positions, the selected procedure showed the lowest mean reconstruction error across deployments; the mean of the deployment-specific median errors was 219 m (Fig. 1**b**; Supplementary Fig. S2). As the GPS positions refer to the accompanying vessel, this comparison supports broad-scale rather than pointwise spatial accuracy.

**Figure 1.**
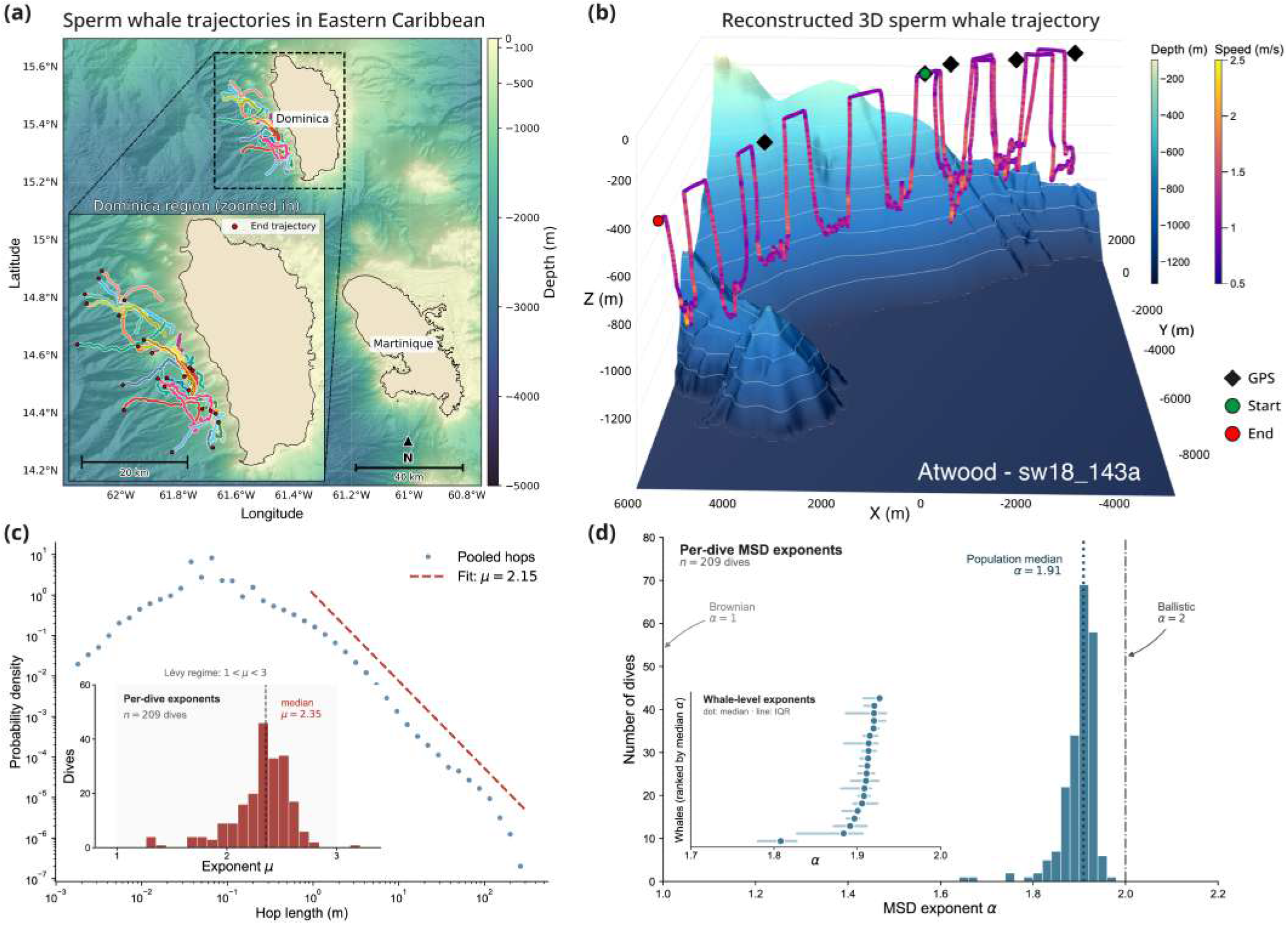
Reconstruction and trajectory-scale organization of sperm-whale foraging movement. **(a)** Reconstructed trajectories from DSWP tag deployments off Dominica, shown over regional bathymetry [1, 2]. The inset shows the spatial distribution of tracks around Dominica; red points mark the end of each reconstructed trajectory. **(b)** Representative three-dimensional trajectory from *Atwood* (deployment sw18_143a), reconstructed using pitch-adaptive dead reckoning and coloured by swim speed. Black diamonds show available vessel-based GPS positions, which provide an independent reference for evaluating broad-scale spatial fidelity. Green and red circles mark the beginning and end of the reconstructed track, respectively, and the underlying surface shows local bathymetry. **(c)** Probability density of step lengths between successive turning points, pooled across *n* = 209 foraging dives. The dashed line shows the fitted power-law tail, with exponent *µ* = 2.15. The inset shows the distribution of exponents fitted separately to each dive, with a median of *µ* = 2.35; most estimates fall within the range conventionally associated with Lévy-like movement (1 *< µ* ≤3). **(d)** Distribution of mean-squared-displacement exponents fitted separately to the *n* = 209 foraging dives. The population median, *α* = 1.91, indicates strongly superdiffusive movement approaching the ballistic limit (*α* = 2), rather than Brownian diffusion (*α* = 1). The inset shows the median and interquartile range of the exponent for each of the *N* = 20 whales, confirming the super-diffusive behaviour at the individual level.

The reconstructed tracks extended from the coastal waters off Dominica into deeper offshore areas over the course of the deployments (Fig. 1**a**). Variations in reconstructed swim speed also delineated the principal phases of the dives, including rapid descent and ascent transits and more tortuous movement at foraging depth. Comparison with bathymetric data [2] showed that foraging occurred predominantly in the open-water pelagic zone, spanning mesopelagic (200–1000 m) and bathypelagic (1000–4000 m) depths. Whales generally remained more than 100 m above the seafloor, and only a small fraction of dives involved movement close to the benthos.

Beyond their spatial layout, the reconstructed trajectories revealed a characteristic statistical organization of search movement. Decomposing the foraging portions of dives into approximately straight segments between successive turning points yielded heavy-tailed step-length distributions (Fig. 1**c**; Supplementary Fig. S3). Estimated power-law exponents were consistent across levels of aggregation: the median dive-level exponent was *µ* = 2.35, the median individual-level exponent was *µ* = 2.36 across the *N* = 20 whales, and the pooled population estimate was *µ* = 2.15 (Supplementary Fig. S4). These values fall within the range conventionally associated with Lévy-like movement (1 *< µ* ≤ 3), in which frequent short displacements are interspersed with rarer long relocations [24–26].

The same scale dependence was evident in the dynamics of displacement. Across dives, the mean-squared displacement scaled as ⟨*r*^2^(*τ*) ⟩ ~ *τ*^*α*^, with a global exponent of *α* = 1.91 (Fig. 1**d**). Individual dives were consistently superdiffusive (*α >* 1), while the scaling exponent decreased with temporal lag: movement was nearly ballistic at lags below 1 min (*α* ≈ 1.98), but progressively less persistent at intermediate (*α* ≈ 1.75, 1–5 min) and longer lags (*α* ≈ 1.52, 5–15 min). The multi-minute estimates use the whole dive excluding the surface phase because the foraging segment alone is too short to support these lags reliably (Supplementary Figs. S5 and S6). Directional persistence was therefore strongest locally but remained detectable over longer portions of a dive. Together, the step-length and displacement statistics identify a multiscale, directionally persistent organization of foraging movement rather than simple diffusive wandering.

We interpret these results as evidence of Lévy-like movement structure, rather than proof that whales implement a unique or optimal Lévy search rule. Similar heavy-tailed and superdiffusive signatures can also emerge from mixtures of behavioural states or interactions with heterogeneous environments [27, 28]. Whales also occasionally returned to previously occupied locations at depth; these “comeback” events are analysed in Supplementary Section III B and Supplementary Figs. S8 and S9.

### Two stereotyped prey-capture tactics identified by acoustic–kinematic clustering

Having characterized how whales organize search over complete foraging trajectories, we next asked whether individual prey-capture attempts are themselves resolved into recurrent acoustic–kinematic tactics. To investigate this question, we analyzed 3,686 prey-capture attempts (averaging 15.62*±*4.87 buzzes per complete dive; 184.3 *±* 135.2 buzzes per individual). Buzzes, defined as rapid sequences of echolocation clicks with inter-click intervals *<* 0.2785 s (sensu [10, 29]), serve as a reliable acoustic proxy for a prey-capture attempt and are acoustically distinct from search-phase echolocation [7]. Figure 2**a** shows the location of distinct acoustic events during an example dive.

Each buzz exhibits a stereotyped internal structure that can be partitioned into three temporal phases based on click-rate dynamics: an initial *ramp-up*, a low-variability *plateau*, and a *final* post-peak decline (Fig. 2b). Details and robustness assessments for this segmentation are provided in the Methods and Supplementary Fig. S10. During the ramp-up phase, click rate increases as inter-click intervals shorten [13, 14]. Ramp-up duration provides a reproducible acoustic measure of how long this acceleration preceding a capture attempt is sustained (Fig. 2c). Together with concurrent movement features, it therefore captures variation in the temporal and kinematic organization of individual buzzes.

**Figure 2.**
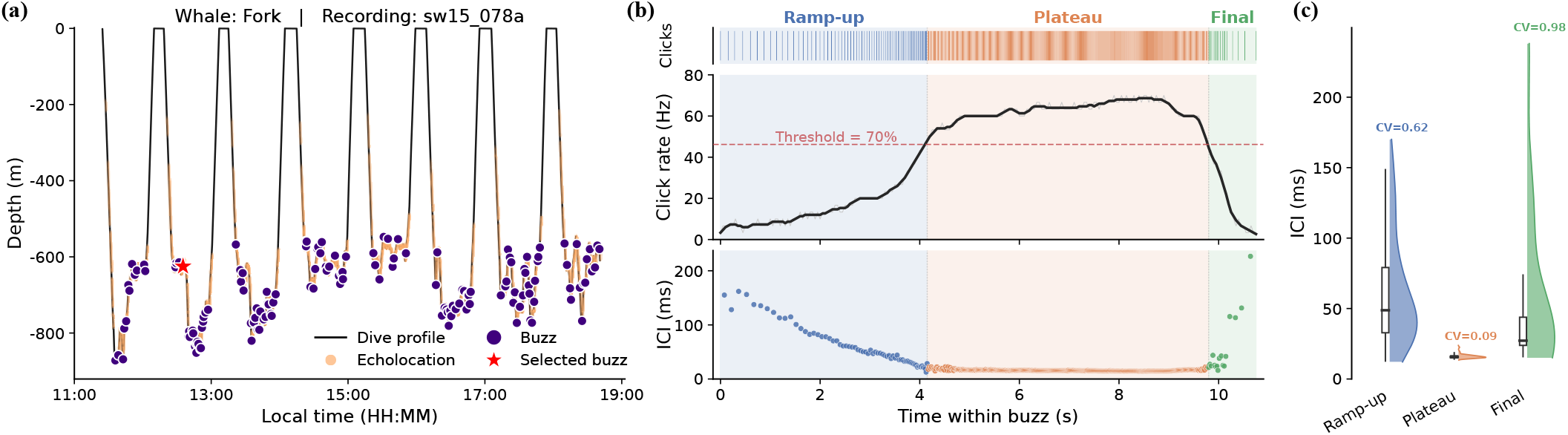
Buzz structure during sperm whale foraging dives. **(a)** Depth profile of a representative recording (whale Fork, sw15_078a), with individual echolocation clicks (orange) and buzz events (purple) overlaid. The starred marker indicates the buzz examined in **(b–c). (b)** Internal structure of a single buzz shown via a click raster (top), instantaneous click rate (middle), and inter-click interval (ICI; bottom), color-coded by phase. Three distinct phases — ramp-up, plateau, and final — delineated using a relative threshold set at 70% of the difference between the maximum and minimum click rates within the buzz (dashed line); the plateau is the high-rate region above this threshold. **(c)** ICI distributions across the three phases confirm the segmentation: the plateau is characterized by markedly lower and less variable ICI (CV = 0.09) relative to the ramp-up (CV = 0.62) and final (CV = 0.98) phases.

Importantly, these acoustic cues coincide with distinct kinematic shifts, such as sudden accelerations or intense maneuvering [13, 30], suggesting a link to different underlying prey-capture tactics. For instance, a representative 100-second 3D trajectory containing two consecutive buzzes illustrates a dynamic shift in behavior within the same dive, as the whale markedly increases its swim speed prior to the second prey-capture buzz (Fig. 3**a**).

**Figure 3.**
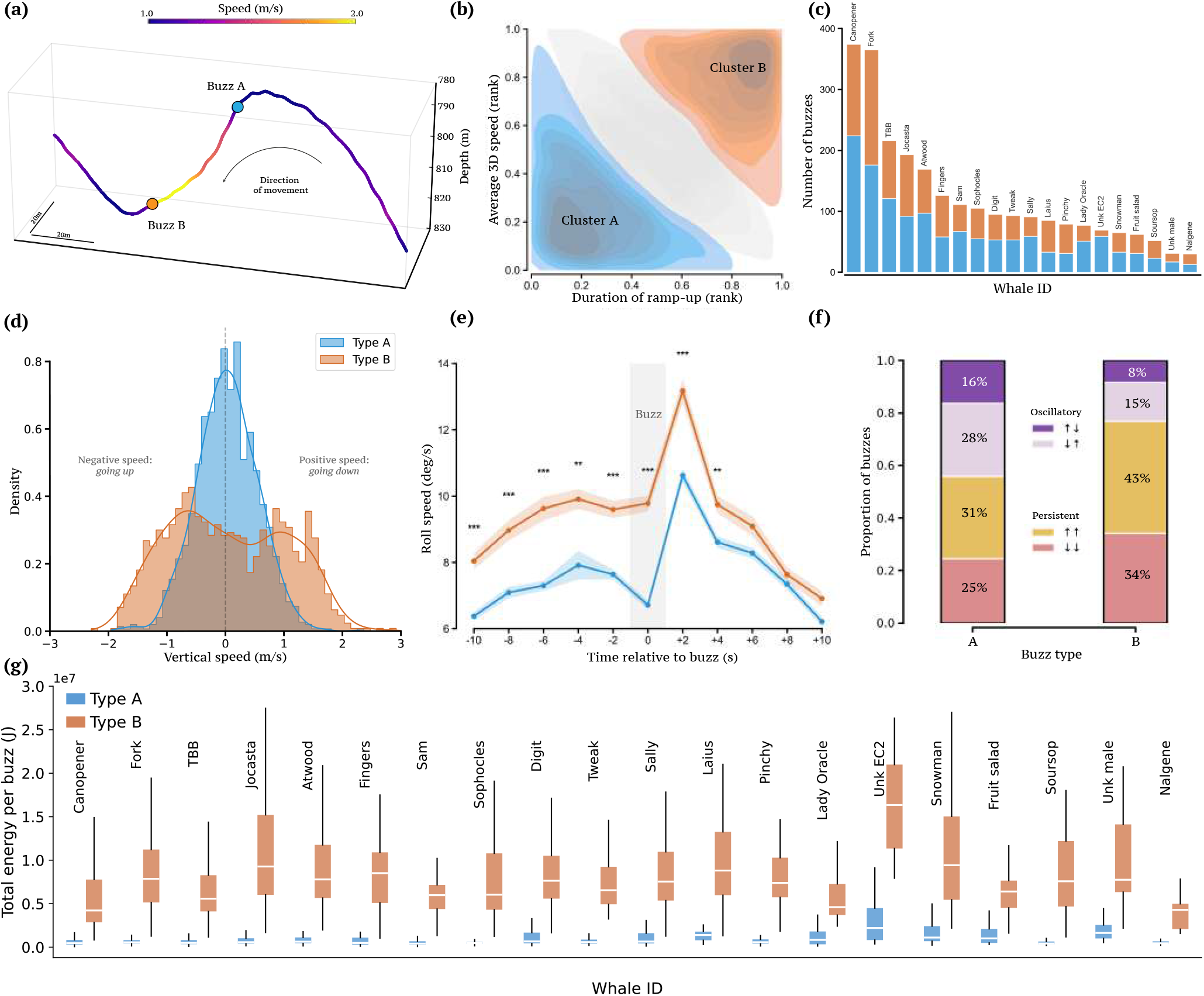
Identification and kinematic signatures of two distinct prey-capture tactics. **(a)** Representative three-dimensional trajectory fragment (duration 100 s) illustrating the occurrence of both buzz types. **(b)** Bivariate kernel density estimate (KDE) of rank-transformed acoustic and kinematic features (*Duration of Ramp-up* vs. *Average 3D Speed*) revealing a bimodal distribution. A Gaussian mixture model identified two distinct clusters: Cluster A (blue; *n* = 1,346), characterized by low swim speeds and brief ramp-up phases; and Cluster B (orange; *n* = 1,142), defined by high-speed pursuits and prolonged ramp-up phases. Buzzes not assigned to either cluster are shown in grey. **(c)** Inter-individual distribution of buzz types across *N* = 20 whales, showing that both tactics occur in every sampled whale. **(d)** Distribution of average vertical speeds in the 10 s preceding buzz onset, revealing a unimodal approach for Type A and a bimodal, high-intensity vertical speed for Type B. Positive values indicate movement toward the seafloor. **(e)** Mean roll speed (deg/s) calculated in bins of 2 seconds centered on the buzz. In the period before the buzz onset, type B events exhibit significantly higher rotational intensity compared to Type A (shaded areas represent standard errors; stars indicate *P <* 0.01, Bonferroni-corrected t-test). **(f)** Proportional distribution of vertical movement classes (oscillatory vs. persistent). Type B attempts show a marked shift toward persistent vertical directionality (77%) compared to the more frequent oscillatory maneuvers in Type A (44%). **(g)** Per-buzz energy estimated by the energetic model across *N* = 20 whales, separated by buzz type. Mean whale-level estimates are 1.28 MJ for Type A and 9.07 MJ for Type B (*P <* 0.001, paired t-test).

To characterize these two distinct prey capture tactics from both an acoustic and kinematic perspective, we extracted a multivariate feature set for each event. This comprised three acoustic features (median click rate, ramp-up duration, and total buzz duration) and two concurrent kinematic features (average 3D swim speed and average roll speed). An initial examination of the pairwise correlations and marginal distributions of these five features revealed a clear dichotomy in the data (Supplementary Fig. S11). This separation was particularly evident in the interaction between one acoustic and one kinematic metric: the duration of the initial acoustic ramp-up and the whale’s average swim speed.

To formally evaluate this dichotomy, the features were first quantile-transformed to approximate marginal Gaussian distributions. This step minimizes the influence of outliers and stabilizes covariance estimation for subsequent clustering. We then examined this joint distribution in rank space—where replacing each feature with its empirical quantile rank decouples the underlying dependence structure from its marginal scales (see Methods). Visualizing the data in this space revealed two well-separated clusters aligned strictly along the axes of ramp-up duration and average swim speed (Fig. 3**b**). Applying unsupervised clustering via a two-component Gaussian Mixture Model (GMM) with full covariance matrices formally separated the data into two clusters representing the two prey-capture tactics (Fig. 3**b**; Supplementary Fig. S12). The optimal number of components was determined by minimizing the Bayesian Information Criterion over *k* ∈ *{*1, …, 8*}*; the minimum was achieved at *k* = 2 (Supplementary Fig. S13). Buzzes with maximum posterior cluster probability below 0.75 were considered unassigned to either cluster and were not used in cluster-specific comparisons (see Methods). This yielded 1,346 Type A and 1,142 Type B buzzes, comprising 54.1% and 45.9% of classified events, respectively. Both tactics occurred in every sampled individual (Fig. 3**c**), showing that the contrast is shared across the 20 whales rather than driven by individual specialization. The propor tion of Type B buzzes within each individual’s classified repertoire ranged from 14.5% to 61.2% (Supplementary Fig. S14). The two tactics also occurred across a similar depth range. Type A buzzes were marginally deeper than Type B (mixed-model estimate +10.6 m, 95% CI 4.5–16.8 m, *P* = 6.5 *×* 10^−4^), but the effect size was negligible (Cliff’s *δ* = 0.08, Cohen’s *d* = 0.13) and the distributions overlapped almost entirely (Supplementary Fig. S15). Thus, depth did not meaningfully distinguish the two tactics.

### Kinematic signatures distinguish prey-capture tactics

Having partitioned the buzzes into two distinct clusters based solely on ramp-up duration and mean swim speed, we next investigated whether they capture broader differences in fine-scale movement dynamics. Specifically, we examined independent kinematic signatures — such as vertical velocity and body roll — in the seconds surrounding buzz onset. Vertical speed distributions in the 10 s window preceding buzz onset reveal a kinematic distinction between the two tactics (Fig. 3**d**). Type A events exhibit a narrow, unimodal distribution tightly centered near zero, showing that these capture attempts occur during near-horizontal, low-speed swimming. In contrast, Type B events show a broader, bimodal distribution with substantially higher absolute vertical speeds (Mann–Whitney *U* test, *P <* 0.001). The two peaks in this distribution correspond to buzzes performed while the whale is moving either steadily downward or upward.

This marked difference in movement activity is also reflected in the whales’ rotational dynamics, aligning with previous studies that link body orientation changes to prey-capture effort [13, 18]. Roll speed, measured in 2-s bins across a 20-s window centered on buzz onset, diverged significantly between the two tactics approximately 8 to 10 s before acoustic onset (Fig. 3**e**). While both types showed increasing roll speed, Type B events exhibited consistently higher roll speeds throughout the pre-buzz and early-buzz periods (Bonferroni-corrected t-test, *P <* 0.01).

To characterize the direction of vertical movement around each buzz, we classified the vertical trajectory in a 10-s window before and after buzz onset by the sign of the vertical velocity, yielding four classes: up–up and down–down (persistent), and up–down and down–up (oscillatory). The class proportions differ between tactics (*χ*^2^ = 122.6, *P <* .001; see Supplementary Fig. S16). Type A buzzes are more variable in direction: 44% are oscillatory (16% up–down, 28% down–up), while the remaining 56% are split between persistent downward (31%) and persistent upward (27%) movement. Type B buzzes, by contrast, are predominantly persistent (77%; 34% downward, 43% upward), indicating a sustained, directed vertical trajectory in a single direction, rather than up-and-down motion.

To test whether these kinematic differences translate into an energetic asymmetry, we used an energy model inspired by previous sperm-whale and cross-species formulations [30, 31] and modified to account for the motor demand represented in the reconstructed trajectories (see Methods). The energetic model integrates work against parasite drag, work during positive tangential acceleration, and resting metabolism over the observed buzz-speed series. Across all twenty individuals, the energy estimated for Type B exceeded Type A by approximately a factor of seven (1.28 MJ vs 9.07 MJ; paired t-test, *P <* 0.001; Fig. 3**g**), with the rank order preserved within every whale (Supplementary Section V A).

Together, these measurements define different estimated energetic investments. Type A buzzes combine lower swim speeds with brief ramp-up phases and lower estimated energy use, whereas Type B buzzes combine higher speeds with prolonged ramp-up phases, greater roll, persistent vertical movement, and higher estimated energy use. At the coarser displacement scale, however, buzz-centred MSD did not distinguish Type A from Type B at any tested lag (Supplementary Fig. S7), indicating that the contrast is concentrated in local kinematics and acoustic timing rather than net displacement around buzz onset.

### Sequential organization and post-buzz acoustic behavior

While our analyses thus far have characterized capture attempts in isolation — establishing two energetically and kinematically distinct tactics (Fig. 3) — the order in which a whale deploys them within a dive offers a complementary, sequence-level view of foraging. To investigate how these buzz types are organized sequentially, we modeled-buzz sequences as a first-order Markov chain, augmenting the buzz-type state space with discrete *Start* and *End* states (Fig. 4**a**). Transition probabilities were estimated as conditional probabilities, and their sensitivity to deployment sampling was quantified across 200 leave-subset-out resamples, randomly excluding ten deployments per iteration (see Methods). Although Type A buzzes were slightly more common than Type B buzzes (*N*_*A*_ = 1346 versus *N*_*B*_ = 1142), both types showed within-tactic persistence. The most likely transition from Type A was to another Type A buzz (*A* → *A* = 0.55 *±* 0.01), whereas Type B buzzes were followed by Type B (*B* → *B* = 0.42*±*0.01). Type B buzzes were also more likely to occur at the beginning of a dive (Start → *B* = 0.54 *±* 0.02; Start → *A* = 0.46 *±* 0.02). Moreover, a pronounced functional asymmetry emerged in the termination dynamics: type B buzzes were twice as likely to precipitate the end of a dive (0.13*±*0.01) compared to type A (0.06*±*0.01). This asymmetry was stable to the removal of sets of deployments. Cross-transitions (A→B: 0.39 *±* 0.01; B→A: 0.45 *±* 0.01) indicate that switching between tactics is frequent, roughly every 2– 3 buzzes on average.

**Figure 4.**
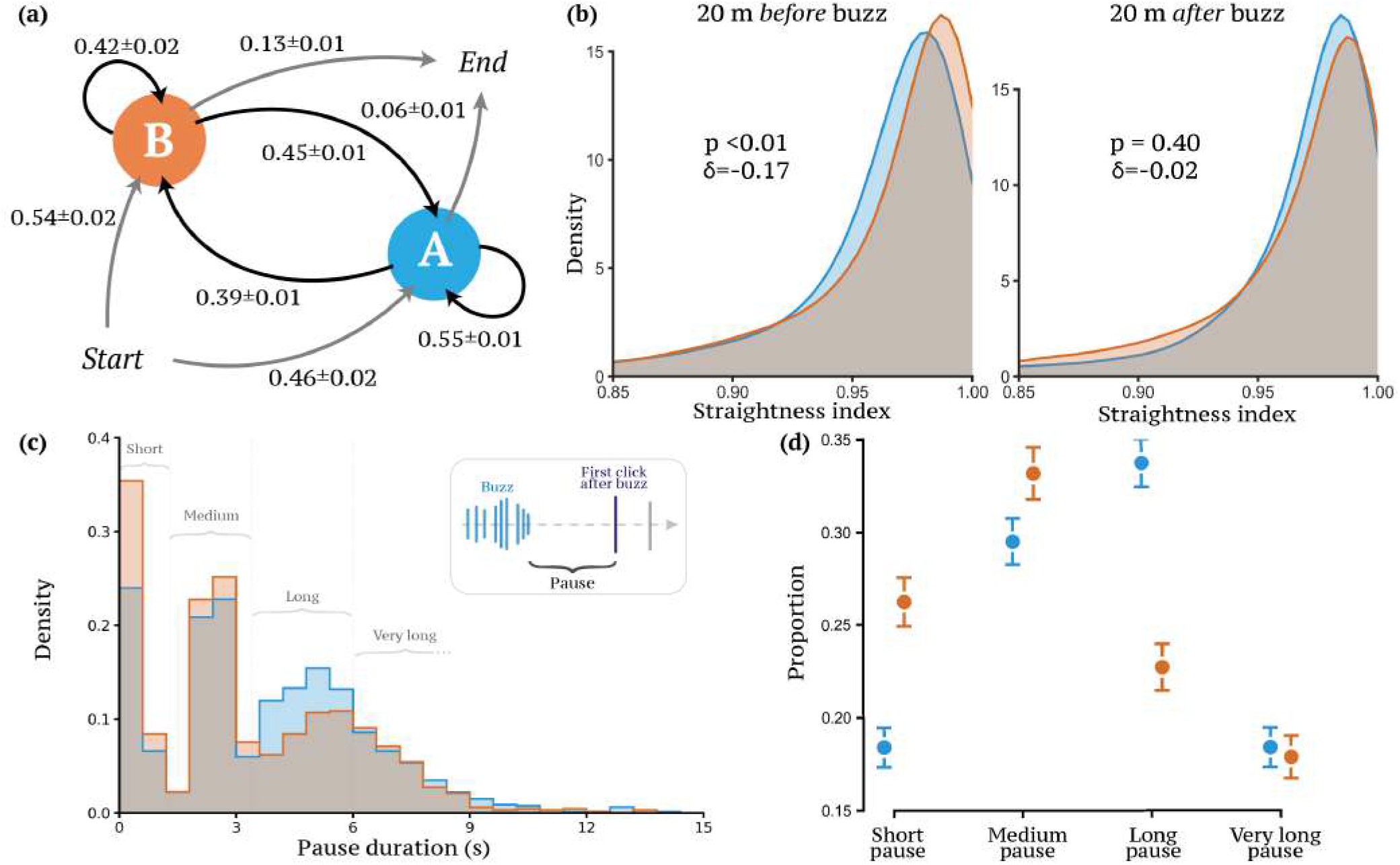
Sequential organization and kinematic consequences of prey-capture tactics. **(a)** State transition diagram representing buzz sequences as a first-order Markov chain. Nodes define behavioral states (*Start, End, Cluster A, Cluster B*), with arrows indicating transition probabilities. Values represent mean *±* s.d. across 200 leave-subset-out resamples. Type B events are twice as likely to terminate a dive (0.13 *±* 0.01) compared with Type A (0.06 *±* 0.01). **(b)** Probability density of the straightness index (ratio of displacement to path length) computed in a 20-m window preceding buzz onset (left) and following the buzz end (right). Type B pursuits exhibit significantly higher pre-buzz linearity (Mann–Whitney *U* test, *P <* 0.01), while post-buzz kinematics are statistically similar between tactics (Mann–Whitney *U* test, *P* = 0.40). **(c)** Normalized probability density of post-buzz pause durations (time until subsequent search click). Vertical boundaries (1.3 s, 3.4 s, 6.0 s) define the four categorical classes used for analysis. **(d)** Proportional distribution of pause categories for Type A (blue) and Type B (orange) buzzes. Error bars represent binomial standard errors. A significant association exists between prey-capture tactic and pause duration (*χ*^2^ = 45.4, *P <* 1 *×* 10^*™*4^, permutation test).

The geometric structure of the approach trajectory reinforces this interpretation. Path tortuosity was quantified using the straightness index (*S* = *D/L*), defined as the ratio between beeline displacement (*D*) and cumulative path length (*L*) computed over a 20-m window [32, 33]. This dimensionless metric ranges from 0 (highly tortuous movement) to 1 (perfectly straight motion) and measures the directional efficiency of movement trajectories. Because *S* depends on the spatial scale over which it is measured, we computed it over a fixed 20-m path window rather than a fixed-time window, which would otherwise confound the metric with the tactics’ different swim speeds. We evaluated *S* over the 20 m of path immediately preceding buzz onset and the 20 m immediately following the last buzz click (the departure; Fig. 4**b**). Both tactics followed near-straight paths overall (*S* ≈ 0.98). During the approach, Type B buzzes were preceded by marginally straighter paths than Type A (Mann–Whitney *U* test, *P <* 0.01; Cliff’s *δ* = −0.17), whereas after the buzz the two tactics were statistically indistinguishable (*P* = 0.40, *δ* = −0.02; see Supplementary Fig. S17 for other windows). The pre-buzz/post buzz contrast in straightness difference is itself informative: it confirms that the kinematic distinction is concentrated in the approach phase and does not reflect a persistent individual disposition toward straight or tortuous movement.

Post-buzz pauses — defined as the time between the last buzz click and the first subsequent search-phase click — further distinguish the two tactics. Final buzzes (those not followed by any further echolocation before surfacing) were first identified and excluded from pause analysis; of the 36 terminal events, 32 were Type B and only 4 were Type A, which is itself strongly consistent with the Markov termination asymmetry (expected ratio under the null hypothesis of equal termination probabilities: ~0.5; observed: 0.89; binomial *P <* 0.001). The full distribution of these post-buzz pauses differed between tactics (Fig. 4**c**, Mann–Whitney *U* and Kolmogorov– Smirnov tests, both *P <* 0.001); summarizing them into four duration classes (short, *<* 1.3 s; medium, 1.3–3.4 s; long, 3.4–6.0 s; very long, *>* 6.0 s) confirmed a significant difference in their proportions (*χ*^2^ = 45.4, *P <* 1 *×* 10^−4^, permutation test, 10,000 iterations; Fig. 4**d**). Type B buzzes were disproportionately followed by short pauses, whereas Type A buzzes were more frequently followed by long or very long pauses. Possible interpretations of this asymmetry are considered in the Discussion.

### Independent Azorean validation links prey-capture tactics to prey-field structure

The prey-capture patterns identified so far rest on a single population (DSWP, Eastern Caribbean) and on inferences about the prey field. To test the generality of the two-tactic framework and anchor it to more direct prey-field observation, we analyzed an independent publicly available dataset from a single juvenile sperm whale off the Azores [12]. Crucially, this dataset includes a real-time echogram generated by the whale’s own biosonar, providing a direct acoustic window into the local prey field during each capture attempt. Over a 22-hour recording period, this whale performed 27 foraging dives and a total of 108 buzzes (median 4 buzzes per dive, range 0–8). To ensure comparability with the foraging context of the DSWP dataset, we focused our analysis on the 11 daytime dives (see Methods for selection criteria). These are consistently deep (622–692 m) and yielded 58 individual buzzes for the analysis. Leveraging the echo-trace analysis previously conducted on this dataset [5, 34], we considered the already-identified individual prey echo streams — continuous sequences of echoes reflecting the same organism across consecutive clicks — to compute three metrics characterizing the local prey field for each capture attempt (see Fig. 5**a** for an illustration): *(i)* the total number of prey echoes within a *±* 10 s window around buzz onset, serving as a proxy for local prey density; *(ii)* the mean absolute depth of the ensonified prey (m), derived from the whale’s instantaneous depth and the echo arrival time; and *(iii)* the nearest-prey range (m), representing the distance to the closest detected organism ahead of the whale at the start of the buzz.

**Figure 5.**
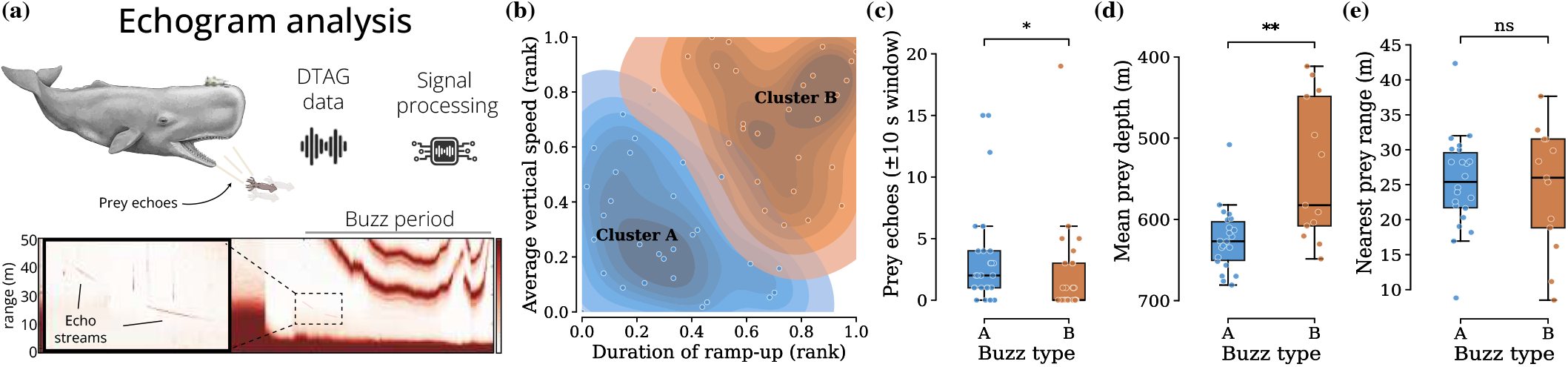
Extreme-corner classification and echogram analysis link prey-capture tactics to distinct prey environments in the Azores. **(a)** Schematic of the echogram analysis based on publicly available Azorean DTAG data [21]. A tag positioned next to a juvenile sperm whale’s blowhole allows for the identification of echo streams from potential prey. Following signal processing [21], the echogram analysis yields target range (m) over time/click indices, identifying individual echo streams and clicks during the buzz period. **(b)** Bivariate KDE of rank-transformed ramp-up duration and average vertical speed for the 58 daytime buzzes (dives 1–11). Type A (blue; *n* = 29) comprises the buzzes closest to the lower-left corner of this feature plane (short ramp-up, low speed); Type B (orange; *n* = 29) comprises those closest to the upper-right corner (long ramp-up, high speed). This 2D extreme-corner classification mirrors the analyses on the DSWP data. **(c)** Number of prey echoes detected within a *±*10 s window around buzz onset. Type A buzzes have a nominally higher echo count (Mann–Whitney *U* test, *P* = 0.037; Cohen’s *d* = 0.35), although this difference is not robust to stricter inclusion criteria or threshold-free analysis. **(d)** Mean prey depth for the two buzz classes. Type A buzzes are associated with significantly deeper prey (*P* = 0.003; Cohen’s *d* = 1.39), consistent with targeting denser prey patches at greater depth during the bottom phase of the dive. **(e)** Nearest-prey range for the two classes, showing no significant difference. These results indicate that the two buzz classes occur in systematically different local prey environments, supporting their interpretation as alternative prey-capture tactics.

Unlike the large, multi-individual DSWP dataset, the smaller Azorean sample (*n* = 58 daytime buzzes) lacked the clear bimodal structure required to support robust unsupervised GMM clustering. Furthermore, because the processed dataset lacked the complete orientation and trajectory variables used to reconstruct DSWP 3D swim speed, we substituted pressure-derived vertical speed — a metric that reproduced the primary GMM classification with high agreement in DSWP (see Supplementary Section IV and Supplementary Figs. S12 and S18). To classify these buzzes, we applied a deterministic, two-dimensional extreme-corner approach in this modified rank-space feature plane. Given the limited sample size, we partitioned the available buzzes based on their proximity to two behavioral extremes. Specifically, the 29 buzzes closest by Euclidean distance to the lower-left corner (short ramp-up, low vertical speed) were classified as Type A, and the remaining 29 buzzes closest to the upper-right corner (long ramp-up, high vertical speed) were classified as Type B (Fig. 5**b**). Unlike our primary DSWP analysis, this balanced bisection categorizes every capture attempt to preserve sample size, inherently retaining intermediate cases that blur the boundary.

Analysis of the two tactics revealed differences in local prey density, with Type A buzzes associated with a higher number of ambient prey echoes in the surrounding *±*10,s window (Fig. 5**c**). However, this difference was not significant under stricter inclusion criteria or in the threshold-free rank test (*ρ* = −0.22, *P* = 0.095). By contrast, the distinction between tactics was tightly and robustly linked to prey depth. Type A buzzes — the short-ramp-up, low-vertical-speed extreme — were associated with significantly deeper mean prey depth than Type B events (Mann–Whitney *U* test, *P* = 0.003; Cohen’s *d* = 1.39; Fig. 5**d**). This depth result persisted across more conservative splits that excluded intermediate buzzes (e.g., *N* = 20, *p* = 0.003; *N* = 25, *p* = 0.001) and was confirmed by a threshold-free Spearman analysis across all 58 buzzes (*ρ* = −0.46, *P* = 0.005). Kinematically, the vertical trajectory around buzzes reproduces the DSWP pattern. Classifying the 10 s before buzz onset and the 10 s after buzz end by the sign of the vertical velocity, Type A buzzes tend to be *oscillatory* (direction reversed between the two windows) and Type B *persistent* (movement sustained in one direction). Under the 29/29 split, this difference is already in the DSWP direction (Type B 61% persistent vs Type A 52%), and it sharpens when the split is restricted to the most extreme buzzes of each tactic, approaching the DSWP values (44% oscillatory for A, 77% persistent for B). Post-buzz pause categorization showed no significant difference between types (*χ*^2^ = 1.07, *P* = 0.784), as expected given the limited sample. Both tactics also showed within-tactic persistence (A→A = 0.55, B→B = 0.57), although with only 11 daytime dives the Markov structure is too sparse for quantitative comparison with Dominica (Supplementary Fig. S19).

Together, these results anchor the A/B distinction to observable variation in the local prey field and show that its principal kinematic structure is recovered at a different site in the North Atlantic, age class, and classification framework.

### Project CETI 2024 deployments reproduce the core kinematic contrast

We further tested the A/B contrast with independent Project CETI Bio-Logger [35] deployments from 2024 off Dominica. Using the same ramp-up-duration *×* speed feature plane and the extreme-corner classification used for the smaller Azorean dataset, we recovered the core behavioral phenotype: Type B buzzes were faster, involved greater roll, followed straighter approaches, and maintained vertical direction more often, whereas Type A buzzes were slower, deeper, and more frequently associated with direction reversals (Supplementary Section VII; Supplementary Figs. S20 and S21). Together with the Azorean analysis, this additional dataset supports the recurrence of the acoustic–kinematic distinction beyond the DSWP discovery sample.

## DISCUSSION

Taken together, the results identify multiscale foraging organization across two nested levels: a dive-level trajectory search process that structures foraging through the water column, and a buzz-level repertoire of prey-capture tactics that whales switch between within individual dives. The first is expressed through persistent, superdiffusive, Lévy-like trajectories with occasional spatial recurrence. The second comprises two sensory– motor, acoustic–kinematic prey-capture tactics that differ in speed, vertical persistence, roll, approach geometry, and model-estimated energetic demand. OFT provides an organizing interpretation for this hierarchy: a generalist predator hunting in a heterogeneous prey field should maintain tactically distinct capture actions within a broader search strategy [15–17]. We treat this framework as an interpretation rather than a demonstrated mechanism.

Multiple foraging tactics have previously been inferred in deep-diving odontocetes, but the structure of the variation has remained unclear. In Norwegian male sperm whales, hidden Markov models identified several buzz types strongly stratified by dive depth, suggesting specialization on vertically partitioned prey assemblages [18]. In contrast, the two tactics identified here co-occur across the same depth range and frequently alternate within single dives, supporting tactical switching within a shared foraging context rather than separation by larger-scale habitat transitions. This contrast may reflect demographic and ecological context: the Dominica population consists primarily of females and immature animals exploiting a diverse tropical cephalopod community [1, 36–39], whereas high-latitude males forage in environments with stronger prey stratification. Similar within-dive tactical variation has been reported in other deep-diving odontocetes [18, 30], supporting the interpretation that fine-grained behavioral switching is a general feature of mesopelagic predation rather than a peculiarity of any one population.

The acoustic ramp-up phase provides a window into this real-time tactical adjustment. During prey interception, whales shorten inter-click intervals to maintain high temporal resolution of target position [13, 14]. The prolonged ramp-ups and sustained high click rates observed in Type B buzzes are consistent with tracking faster or more evasive prey during extended pursuits, whereas the short ramp-ups and near-zero pre-buzz vertical speeds of Type A are consistent with encounters requiring less tracking effort, for example with slower prey or schooling prey that require less pursuit effort. That fewer than 10% of acoustically detected organisms are ultimately pursued [12] further indicates an active selection process: whales are not merely tracking what is present, but choosing which prey to pursue and how to engage them. Determining whether capture success differs between tactics will require event-level resolution of post-buzz outcomes, for which animal-borne camera tags offer a promising approach [40].

The energetic model separates the two tactics by approximately a factor of seven (1.28 MJ vs 9.07 MJ per buzz across *N* = 20 whales), with Type A lower than Type B in every animal (Supplementary Section V A). These values should be read comparatively: the key result is the marked ratio between the tactics, and its consistency across whales, rather than the exact energy assigned to either tactic. The faster and longer Type B buzzes therefore carry substantially greater model-estimated energetic demand. Read against OFT predictions, this is consistent with a repertoire in which a lower-demand tactic dominates by number while a higher-demand tactic is deployed when its expected return justifies the additional effort [15–17]. The 2*×* higher dive-termination probability of Type B is also compatible with an energetic-budget interpretation, although neither energetic return nor capture outcome is observed directly. The longer post-buzz pauses following Type A (Fig. 4**c,d**) — potentially reflecting prey handling [7] — and the rapid acoustic re-engagement following Type B provide testable predictions. Animal-borne cameras or stomach-contents data could resolve whether Type A and Type B target taxonomically distinct prey, differ primarily in encounter geometry, or differ in capture success.

The recurrence of the same behavioral contrast in two additional datasets supports the interpretation that the A/B distinction reflects a reproducible prey-capture phenotype rather than a peculiarity of one classifier or deployment set. The Azorean data connect this contrast to measured prey-field structure, while the 2024 CETI deployments recover its principal kinematic expression without concurrent echograms. The latter sample remains modest (231 kinematic buzzes from 10 tags and 19 dives; heading-dependent metrics for 149 buzzes from five tags), limiting claims about prevalence and individual consistency, but the positive kinematic convergence supports the recurrence of the core tactic phenotype. Because the two regions host distinct cultural clans and partially different prey assemblages [36–39, 41, 42], this convergence raises the hypothesis that the tactics respond to general prey properties—such as size, trophic position, or whether prey are encountered singly or in aggregations— rather than mapping to particular prey species [43]. Direct prey identification will be needed to test this interpretation.

Beyond any single buzz, the temporal organization of capture attempts reveals additional structure within the foraging dives of sperm whales in Dominica. Within-tactic self-transition probabilities (A→A = 0.55, B→B = 0.42) indicate that once a whale engages a tactic, it often persists across several consecutive attempts before switching. Cross-transitions are not negligible (A→B = 0.39, B→A = 0.45), suggesting tactical reassessment every two to three buzzes on average. Type B also has a 2*×* higher dive-termination probability than Type A. This sequence structure is consistent with cost-balanced organization of foraging bouts. Further data coupling prey-field abundance and species availability with acoustic– kinematic measurements are needed to establish the underlying decision mechanism.

More broadly, the within-dive switching, the energetic asymmetry, and the structured sequence dynamics implicate a flexible decision-making process driven by realtime acoustic feedback. This places sperm whale foraging within the wider work on cognitive flexibility in cetaceans [44, 45] and connects, perhaps unexpectedly, to recent results from the same population showing socially learned vocal structure across cultural barriers [46, 47]: the same individuals that adjust their hunting tactics to local prey on the timescale of seconds also acquire communication-system features on the timescale of years. Whether tactical flexibility and social-learning capacity are mechanistically linked — both expressions of a general cognitive architecture — or instead independent traits that happen to co-occur in this species remains an open question. The buzz-level decision substrate identified here provides a tractable behavioural assay for the first of these capacities; the coda system [46] provides one for the second. Combining the two in the same individuals offers a path toward addressing the question empirically.

The differing maneuvering demands of the two tactics carry a clear practical implication: anthropogenic noise exposure may disrupt them asymmetrically. High-investment Type B pursuits, which combine intense rotational dynamics, prolonged sensory tracking, and greater inferred energetic demand, may be more sensitive to interruption than lower-investment Type A captures within prey patches. This prediction is testable through controlled exposure experiments [48, 49] or opportunistic monitoring during shipping- or military-noise events [50]. Such asymmetric sensitivity has broader conservation implications: aggregate metrics of “foraging activity,” which treat all buzzes as equivalent, may underestimate disruption of high-demand tactics. More broadly, distinguishing tactics makes explicit the difference between what a whale does and what an anthropocentric observer recognizes, an instance of the *umwelt* argument [51] that behavior is best understood within an organism’s perceptual frame. Linking specific echolocation patterns to prey-capture tactics recovers behavioral decisions that would otherwise remain hidden to human observers.

Several limitations qualify these results. First, females and immature whales dominate the dataset; adult males, whose foraging behaviour has been characterised separately [10, 18, 19] and which may exploit different prey assemblages, are absent. Second, dietary inference rests entirely on prey-field context (Azorean echograms) rather than direct identification; stomach-contents data or animal-borne camera footage would be needed to confirm the prey-identity interpretation of the A/B contrast. Third, the prey-field validation rests on a single Azorean deployment, while the independent CETI 2024 validation lacks concurrent echograms and individual demographic metadata; additional individuals and ocean basins are needed to establish generality. Finally, the cost–benefit interpretation of the post-buzz pause asymmetry rests on the assumption that long pauses correspond to prey handling [7] and short pauses to rapid re-engagement; direct verification — for example via acoustic detection of mastication sounds — has been considered in the context of this dataset but is not yet conclusive.

Several concrete next steps would directly test the framework developed here. First, combining sensor-tag audio with animal-borne cameras would directly identify the prey species engaged by each buzz type, resolving the A/B prey-identity question that current data only address indirectly. Second, extending the two-tactic analysis to other sperm-whale populations with characterized prey communities (Norwegian Sea, Gulf of California, Sri Lanka) would test the geographic generality of the framework. Third, applying the same acoustic-kinematic methodology to other deep-diving odontocetes (Cuvier’s beaked whales, pilot whales, narwhals) would reveal whether tactical switching of this kind is a general solution to mesopelagic predation. Fourth, the asymmetric-noise-sensitivity prediction articulated above is directly testable with existing controlled-exposure datasets [49] and would translate this acoustic-behavioural taxonomy into a conservation tool. Finally, coupling buzz-type sequences to the coda communication system [46, 47] in the same individuals would test whether social context predicts tactical choice — a question that requires datasets in which both acoustic streams are recorded synchronously, but which the same DSWP corpus is well positioned to answer.

In conclusion, sperm-whale foraging combines persistent, deliberate search across complete dives with flexible sensory–motor choice between at least two prey-capture tactics. This nested organization is consistent with OFT expectations for an informed predator that adjusts its investment to local encounter conditions [11]. Whether prey information is shared socially, or whether tactic use varies among individuals, social units and vocal clans, remains open. Nevertheless, the recovery of the core acoustic–kinematic contrast across regions suggests that part of this repertoire reflects general constraints of finding and capturing prey in heterogeneous pelagic environments, rather than a strategy specific to a single populartion or prey assemblage.

## METHODS

### Data description

We analysed one primary discovery dataset and two independent validation datasets.

#### DSWP discovery dataset

The primary dataset was obtained from the Dominica Sperm Whale Project (DSWP) and comprised 34 DTAG deployments conducted off the coast of Dominica between 2014 and 2018 [20]. The tags recorded depth from hydrostatic pressure, three-axis acceleration and magnetometry at a common sensor sampling rate of 25 Hz, together with broadband audio at 120 kHz. The sensor and audio streams shared a common tag clock. Orientation (pitch, roll and heading) was derived from the accelerometer and magnetometer measurements and corrected from the tag reference frame to the whale body frame. The dataset spans 268 h of recordings and contains 245 foraging dives deeper than 300 m from 20 individuals, predominantly females and immature animals from the resident Eastern Caribbean population [1]. Buzzes were identified using the Dominica criterion reported by Tønnesen *et al*. [29], as click sequences with inter-click intervals below 0.2785 s.

#### Azorean validation dataset

The first validation dataset was a publicly available 22-h DTAG deployment from a single juvenile sperm whale off the Azores (tag `sw17_196a`) [12]. The deployment comprised 27 foraging dives and 108 buzzes. The tag was positioned near the blowhole and provided hydrostatic-pressure measurements at 25 Hz together with acoustic recordings from which forward-facing prey-field echograms were derived. The available echogram variables included prey-echo count, mean prey depth and nearest-prey range around each buzz. Buzzes in this dataset were defined by inter-click intervals below 0.154 s. The validation analysis used the 11 daytime dives, comprising 58 buzzes, for which the diving context was most comparable to the DSWP dataset. The processed Azorean data did not contain the complete orientation and trajectory variables used for the DSWP three-dimensional speed reconstruction, so kinematic analyses used vertical speed derived from the pressure record.

#### Project CETI 2024 Dominica validation dataset

The second validation dataset comprised independent deployments of CETI Bio-Loggers [35] conducted off Dominica in 2024 by Project CETI. Combined per-tag structures provided dive boundaries, echolocation events and movement measurements, including depth, pitch and roll sampled at 25 Hz. Heading was available for five of the ten tags. Buzzes were detected using the same inter-click interval threshold of 0.2785 s, with a minimum of 30 clicks per event. The kinematic cohort comprised 231 buzzes from 10 tags and 19 dives; 149 buzzes from five tags had valid heading and could therefore be used for heading-dependent path-geometry analyses. The available structures did not contain prey-field echograms or usable whale-identity, age or sex fields. Consequently, this dataset supports an independent test of the kinematic contrast between prey-capture tactics, but not prey-field or per-individual stratified analyses.

Dataset-specific preprocessing, classification and validation procedures are described in the corresponding Methods subsections below.

### Reconstruction of dive trajectories

Fine-scale three-dimensional movement trajectories were reconstructed from the 25-Hz pitch, roll, and heading (yaw) time series recorded by DTAG sensors [20], using a dead-reckoning (DR) framework in which the instantaneous velocity vector is integrated forward in time to produce a cumulative position track. Prior to integration, depth, pitch, and heading signals were smoothed with a 5-sample centered moving-average filter to reduce high-frequency sensor noise. Swimming speed at each time step was estimated using a pitch-adaptive procedure. That is, when the whale was actively ascending or descending — defined by the simultaneous conditions |Δ*d*(*t*)| ≥ *δ*_*z*_ and |*θ*(*t*)| ≥ *θ*_*p*_, where *δ*_*z*_ = 0.05 m sample^−1^ and *θ*_*p*_ = 0.05 rad are depth-change and pitch thresholds, respectively — instantaneous speed was derived from the rate of depth change and the pitch angle:

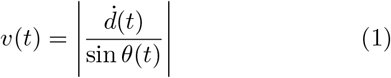

where 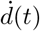 is the numerical depth derivative. The dual condition prevents division by near-zero sin *θ* values that can arise from sensor noise during nominally steep segments. Speed estimates were clipped to [0.1, 5.0] m,s^−1^ to exclude physically implausible values. During near-horizontal segments where either condition was not met, speed was set to a constant fallback value *v*_0_ = 0.9 m,s^−1^, consistent with published mean foraging-dive speeds for sperm whales [3, 52]. Horizontal position was integrated as

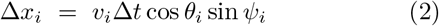

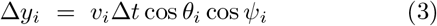

where *ψ*_*i*_is the heading and Δ*t* = 1*/f*_*s*_ = 0.04 s. Depth was set directly from the smoothed pressure sensor record rather than integrated, eliminating vertical drift.

As DTAG units lack internal GPS, trajectories were computed in a local reference frame with origin at the point of tag attachment. The sensor orientations were first corrected from the tag frame to the whale body frame. A further horizontal rotation *α* was then applied to align the local track with geographic coordinates. For each deployment, *α* was selected by a greedy search over {0°, 10°, …, 350°}, retaining the value that minimized the mean nearest-neighbour Euclidean distance between the reconstructed track and vessel GPS positions collected while following the surfaced whale.

A full evaluation of alternative dead-reckoning approaches — including a depth-rate method, a constant-speed model, and an observed climb/descent rate (OCDR) method [7, 23] — and their GPS-error statistics across all deployments is provided in Supplementary Information (Supplementary Section II and Fig. S2).

### Plateau phase detection within buzzes

Each buzz was segmented into three temporal phases — ramp-up, plateau, and final — based on the instantaneous click rate computed from the click timestamps. Click rate was estimated using a sliding symmetric window of duration *w* = 0.3 s, stepped at 0.05 s intervals, yielding a discrete click-rate time series *r*(*t*) (clicks s^−1^) at each window centre *t*. To reduce high-frequency noise, *r*(*t*) was smoothed with a 5-point centered moving average, producing 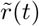.

The plateau was identified as the contiguous temporal region in which 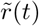 exceeded a relative threshold *τ*, defined as:

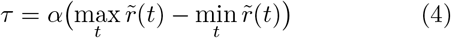

where *α* = 0.7 is a dimensionless scaling parameter. This formulation scales the threshold to the within-buzz click-rate range, reducing sensitivity to differences in scale across individuals and recording conditions. The plateau boundaries [*t*_start_, *t*_end_] were set to the onset and offset of the longest contiguous interval satisfying 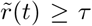. The ramp-up phase spans [*t*_0_, *t*_start_) and the final phase spans 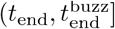, where *t*_0_ and 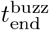 are the times of the first and last click in the buzz, respectively.

The choice of *α* = 0.7 was validated through a sensitivity analysis across *α* ∈ [0.05, 0.95] applied to all *N* = 3,686 buzzes (Supplementary Fig. S10). Two complementary criteria were evaluated. First, the coefficient of variation (CV) of inter-click intervals within the detected plateau decreased with *α*, while its derivative approached a plateau near *α* = 0.7, indicating diminishing gains in regularity at stricter thresholds. Second, each detected plateau was compared with 20 randomly sampled, same-duration windows from the same buzz. Excess regularity, defined as the median null-window CV minus the plateau CV, peaked near *α* = 0.7. Together, these tests indicate that *α* = 0.7 identifies a low-variability plateau (median CV = 0.12) that is maximally distinct from arbitrary segments of the same buzz and robust to small changes in threshold.

### Buzz types from clustering

To investigate the existence of stereotypical foraging behaviors, we analyzed a range of buzz-related features, ranging from acoustic properties, which characterize the click train structure, to behavioral metrics describing whale movement. The feature set included:

- *Median click rate:* The median repetition rate (clicks s^−1^) maintained during the buzz.
- *Duration of ramp-up phase:* The duration of the initial high-acceleration component of the buzz, defined as the phase characterized by a rapid, monotonic increase in click rate.
- *Total duration:* Duration of the full buzz event.
- *Average speed:* The average magnitude of the 3D velocity vector during the buzz.
- *Average roll speed:* The average rotational speed around the longitudinal axis.

Roll speed was expressed in degrees per second; legacy mean absolute roll increments per 25-Hz sample were converted to rates by multiplying by the sampling frequency. We examined pairwise feature relationships by visualizing their joint distributions in rank space. This approach highlights the dependence structure between variables independent of their marginal distributions (see Supplementary Figure S11).

An interesting pattern emerged from the combination of *Duration of ramp-up* and *Average speed*. As illustrated in Figure 3**b**, the bivariate KDE reveals a clear dichotomy in the data structure.

To formally distinguish these clusters, we employed a Gaussian mixture model (GMM). Prior to fitting, each feature was mapped through a normal-score quantile transform to approximate a standard-normal marginal distribution, reducing the influence of outliers and stabilizing covariance estimation. This differs from the empirical ranks rescaled to [0, 1] used only for visualization and extreme-corner classification. We fitted a two-component GMM with full covariance matrices, initialized via K-means, to the 3,488 buzzes with complete ramp-up-duration and average-speed measurements; 198 of the 3,686 total buzzes lacked one of these clustering features and were not entered into the model. For each included buzz, the model estimated its probability of belonging to each cluster. Buzzes were assigned to a cluster when their maximum estimated membership probability was at least 0.75; the 1,000 events below this threshold were classified as ambiguous and excluded from cluster-specific comparisons.

This probabilistic approach identified two distinct clusters (Figure 3**b**):

- **Cluster A (Blue):** Characterized by low-speed events with brief ramp-up phases (lower-left quadrant).
- **Cluster B (Orange):** Defined by high-speed pursuits coupled with prolonged ramp-up phases (upper-right quadrant).

We subsequently analyzed the distribution of these clusters across individuals (Fig. 3**c**) to determine whether they represented idiosyncratic behaviors or a shared repertoire. Both buzz types were identified in every sampled individual. Their frequencies varied, with Type A generally more common and Type B a widespread but less frequent higher-investment alternative.

Figure 2**b** illustrates the click-rate segmentation for a representative buzz. Type-specific inference is therefore based on the measured ramp-up duration and kinematics rather than on this single illustrative time series.

### Depth comparison of buzz types

To test whether Type A and Type B buzzes occurred at different depths while accounting for the non-independence of buzzes recorded within the same dive and the same individual, we fitted a linear mixed-effects model to the mean depth of each buzz (*n* = 2488 classified buzzes: 1346 Type A, 1142 Type B, from 226 dives, 32 deployments and 20 individuals). Buzz type (A vs. B) was entered as a binary fixed effect, and the model included random intercepts for individual whale and for dive nested within whale. Models were fitted by restricted maximum likelihood (REML) using `statsmodels` (v0.14.4, L-BFGS optimiser); the fixed effect is reported as the estimated A−B depth difference with its Wald 95% confidence interval and *p*-value. Effect sizes were quantified with Cliff’s *δ* and Cohen’s *d*.

### Energy-expenditure estimation

Per-buzz energy was estimated with a trajectory-driven time-series model inspired by the steady-swimming formulation of Aoki et al. [30] and the hydrodynamic framework of Goldbogen et al. [31], and modified to account for motor demand using the reconstructed trajectories. The energetic model combines parasite-drag work, work during positive tangential acceleration, and baseline rest-of-body metabolism. Body mass and length were fixed at *M*_body_ = 4.0 *×* 10^4^ kg and *L*_body_ = 12.0 m. For each buzz, the 3D swim-speed series obtained from the pitch-adaptive reconstructed trajectory was supplied over the full interval from the first to the last buzz click. Using the observed speed series in place of an analytical prey-interception trajectory tailors the calculation to the measured buzz kinematics. We assessed sensitivity to model formulation by comparing the energetic model with the steady-drag model of Aoki et al. [30] over the same observed intervals. Full equations, assumptions, component contributions, and comparison results are provided in Supplementary Section V A. For whales with multiple recordings, Type A and Type B energy estimates were pooled across recordings before the resulting individual-level means were compared using a paired t-test. Because both buzz types use identical body-morphology parameters within each whale, the A-vs-B comparison and its per-individual rank order are robust to absolute body-mass mis-specification; only absolute energy values would shift. Full equations, parameter table, and the implementation code are provided in SI *§* Energy-expenditure model.

### Dynamics across buzz phases

The structure of a movement path reflects the underlying behavioral processes that produced it [22]. We quantified this geometric structure through tortuosity metrics. The Straightness Index, as defined by Batschelet (1981) [32], provides a single summary measure by calculating the ratio of the beeline distance between the trajectory start and endpoint to the total distance traveled along the path:

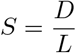

where *D* is the straight-line distance between the first and last position, and *L* is the cumulative path length. Values range from 0 (maximally tortuous) to 1 (perfectly straight). Differences in the distribution of *S* between buzz types were assessed using a Mann–Whitney *U* test, with effect size reported as Cliff’s *δ*.

### Post-buzz pause differences

The post-buzz pause was defined as the time interval between the last click of a buzz and the first click of the subsequent echolocation train. Buzzes that occurred at the end of the vocal phase (terminal buzzes) were identified by the absence of subsequent echolocation clicks before surfacing and excluded from the pause-duration analysis. The continuous pause distributions were compared between Type A and Type B with a two-sided Mann–Whitney *U* test. To localize where the distributions diverged, durations were divided into four non-overlapping classes: Short (*<* 1.3 s), Medium (1.3 ≤ *t <* 3.4 s), Long (3.4 ≤ *t* ≤ 6.0 s), and Very Long (*>* 6.0 s). Association between tactic and pause class was tested with a *χ*^2^ test of independence on the resulting 2*×*4 contingency table and confirmed by permutation (10,000 label shuffles).

### Markov modeling of buzz type transitions

We investigated the temporal organization of foraging events by modeling the sequence of buzzes within a single dive as a discrete-time, first-order Markov chain. The state space defined by the two buzz clusters (*S* ∈ *{A, B}*) was augmented with abstract ‘Start’ and ‘End’ states to quantify the likelihood of each type initiating or terminating a sequence. Transition probabilities were calculated as the maximum-likelihood estimate of *P*_*ij*_ = *P*(*S*_*t*+1_ = *j* | *S*_*t*_ = *i*). Ambiguous buzzes were retained as gaps in the ordered sequence: transitions touching an ambiguous event were discarded, rather than bridging the classified events on either side. To assess sensitivity to highly sampled deployments, the transition matrix was recomputed 200 times after randomly excluding ten deployments in each iteration. The figure reports the mean probability and standard deviation across these resamples.

### Azorean buzz classification

To further validate our approach, we turn to a different data source and apply our methodology. The Azorean dataset is a publicly available DTAG deployment on a single juvenile sperm whale (tag ID sw17_196a) [12]. In this dataset, buzzes were distinguished by inter-click intervals below 0.154 s. The DTAG was mounted near the blowhole, providing a forward-facing echogram of the prey field during foraging dives. We used the available echogram statistics and aggregated across specific time intervals: prey echo count in a *±*10 s window centered on each buzz, mean prey depth, and nearest-prey range. Vertical speed was derived from the 25 Hz hydrostatic pressure sensor as the mean signed depth derivative (m/s) over the buzz interval, smoothed with a 3-point boxcar kernel (positive = descending).

The dataset contains 27 foraging dives; dives 1–11 were classified as daytime (local sunrise to sunset) and are consistently deep (622–692 m), comparable to the DSWP mesopelagic foraging context. Dives 12–27 include progressively shallower and nocturnal events and were excluded to ensure a comparable foraging context. This yielded *n* = 58 buzzes from the daytime subset.

Unlike the multi-individual DSWP dataset, the smaller Azorean sample (*n* = 58 daytime buzzes) lacked sufficient structural bimodality to support robust unsupervised GMM clustering. Because the processed dataset lacked the complete orientation and trajectory variables used to reconstruct DSWP 3D swim speed, we substituted pressure-derived vertical speed, a metric that reproduced the primary classification with high agreement in DSWP (Supplementary Section IV). To classify the Azorean buzzes, we applied a deterministic, two-dimensional extreme-corner classification in rank space (ramp-up-duration rank *×* vertical-speed rank, both rescaled to [0, 1]). Ramp-up duration was computed using parameters identical to the DSWP analysis. Type A was defined as the *N* buzzes with the smallest Euclidean distance to the lower-left corner (0, 0), and Type B as the *N* buzzes closest to the upper-right corner (1, 1). Setting *N* = 29 classified all 58 daytime buzzes with no overlap or unclassified events. Analyses were repeated using more conservative split sizes (*N* = 20 and *N* = 25), which excluded intermediate cases. Between-group comparisons used two-tailed Mann–Whitney *U* tests, with effect sizes reported as Cohen’s *d*. A continuous, threshold-free validation used Spearman’s rank correlation between the diagonal rank score and mean prey depth across the unpartitioned dataset.

### Project CETI 2024 validation dataset and classification

The independent Project CETI 2024 validation used combined per-tag structures from Dominica deployments containing movement, dive, echolocation, and metadata fields. Movement variables (depth, pitch, roll, and, where available, heading) were sampled at 25 Hz. Buzzes were detected using the DSWP criterion (inter-click interval *<* 0.2785 s and at least 30 clicks), and three-dimensional trajectories were reconstructed per dive using the validated pitch-adaptive dead-reckoning implementation. The kinematic cohort comprised 231 buzzes from 10 tags and 19 dives. Heading was available for five tags, yielding 149 buzzes for horizontal path-geometry metrics; swim-speed magnitude remained available for all 231 kinematic buzzes because it is independent of heading orientation.

We used the extreme-corner classifier to test the behavioral contrast in the ramp-up-duration *×* 3D-speed feature plane. After quantile transformation and rescaling both axes to [0, 1], Type A comprised the 115 buzzes nearest the lower-left corner (short ramp-up, slow movement) and Type B the 106 buzzes nearest the upper-right corner (long ramp-up, fast movement). Between-tactic comparisons used two-sided Mann–Whitney *U* tests with Cliff’s *δ* for continuous features and Fisher’s exact test for oscillatory versus persistent vertical geometry. The available structures did not include prey echograms or usable whale identity, age, or sex fields. Full definitions, figures, and caveats are provided in Supplementary Section VII.

### Statistical analysis

All tests were two-tailed. Multiple-comparison corrections were applied using the Bonferroni procedure where indicated. Permutation-based significance thresholds were computed from 10,000 random label permutations. All analyses were implemented in Python (3.9+) using NumPy, SciPy, and scikit-learn.

## DATA AND CODE AVAILABILITY

Analysis code and publication-safe derived data will be deposited in a public repository before publication, with the permanent URL added here. The release contains pseudonymized buzz-level measurements and reconstructed trajectories in local coordinates, but not raw DTAG audio, raw sensor streams, absolute GPS positions, field identifiers, or identifier crosswalks. The source data for the Azorean validation are openly available from Dryad at https://doi.org/10.5061/dryad.tqjq2bvwg.

## AUTHOR CONTRIBUTIONS

L.Z. and C.M. contributed equally to this work. G.P., A.S., and C.M. conceptualized the study. L.Z., C.M., and A.S. led the main data analysis. L.Z. developed software, curated data, performed formal analyses, and prepared visualizations. C.M. developed the clustering and sequence-analysis methodology, performed formal analyses, prepared visualizations, and contributed supervision. S.P. performed the data analysis of the Lévy-walk component. M.L. contributed acoustic analysis and interpretation, sequence interpretation, and conceptual framing. H.B. performed the analysis of trajectory properties. R.D. developed and performed the energetic analysis and curated the associated data. P.T. contributed data and resources, validation, and biological interpretation. D.G. contributed Project CETI resources, funding acquisition, project administration, supervision, and conceptual framing. S.G. contributed data and resources from the Dominica Sperm Whale Project, field investigation, project administration, and biological interpretation. A.S. led trajectory reconstruction and computational and methodological development, and contributed software, formal analysis, validation, visualization, project administration, and supervision. G.P. supervised and administered the project, acquired funding, and led manuscript preparation. All authors contributed to the interpretation of the results, reviewed and edited the manuscript, and approved the final version.

## COMPETING INTERESTS

The authors declare no competing interests.

## ACKNOWLEDGEMENTS

G. P. is supported by the European Research Council (ERC) Consolidator Grant under the European Union’s Horizon Europe programme (grant agreement No. 101171380, project RUNES), and “BeyondTheEdge: Higher-Order Networks and Dynamic” (REA Grant Agreement No. 101120085). We are grateful to Cláudia Oliveira, Mark Johnson and Peter Teglberg Madsen for making the Azores tag data publicly available.

## SUPPLEMENTARY MATERIAL

## I. DATASETS, DEPLOYMENTS, AND ANALYSIS COHORTS

## II. THREE-DIMENSIONAL TRAJECTORY RECONSTRUCTION AND VALIDATION

We reconstructed the three-dimensional (3D) underwater trajectories of sperm whales from depth, pitch, roll, and heading time-series recorded by DTAG biologgers (sampling rate *f*_*s*_ = 25 Hz). All methods follow the dead-reckoning (DR) framework: the instantaneous velocity vector is integrated forward in time to produce a cumulative position track. Because dead-reckoning accumulates heading and speed errors over time, we evaluated four methods that differ in how swimming speed and direction is estimated. To quantify the accuracy of the 3D reconstructions, we rely on the vessel-based GPS points collected concurrently with each deployment, which maintained close proximity to the surfaced whale throughout each recording. For each GPS point (*x*_*g*_, *y*_*g*_) projected into the local Cartesian frame, we computed the minimum 2D Euclidean distance to any point on the reconstructed trajectory:

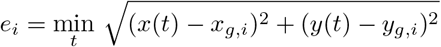

This nearest-neighbour approach avoids time-synchronization issues between the GPS and the DTAG clock. Note that the vessel GPS position represents the ship’s location rather than the whale’s exact position; reported errors are therefore upper bounds on reconstruction inaccuracy. Summary statistics (mean, median, IQR) were computed across all GPS fixes for each recording, and the method with the lowest mean error per recording was designated the “winner” for that deployment.

### A. Coordinate system and sensor conventions

The tag body frame uses a right-handed coordinate system with *x* pointing forward along the whale’s longitudinal axis, *y* pointing to the right, and *z* pointing downward. At each time step *t* the rotation matrix

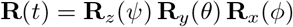

is constructed from heading *ψ*, pitch *θ* (positive nose-up), and roll *ϕ* (positive right-side up) using the ZYX Euler convention. The global frame has *x* pointing North, *y* pointing East, and *z* pointing downward (positive depth). Depth *d*(*t*) is obtained directly from the pressure sensor (accuracy *±*0.1 m) after calibration.

**Figure S1.**
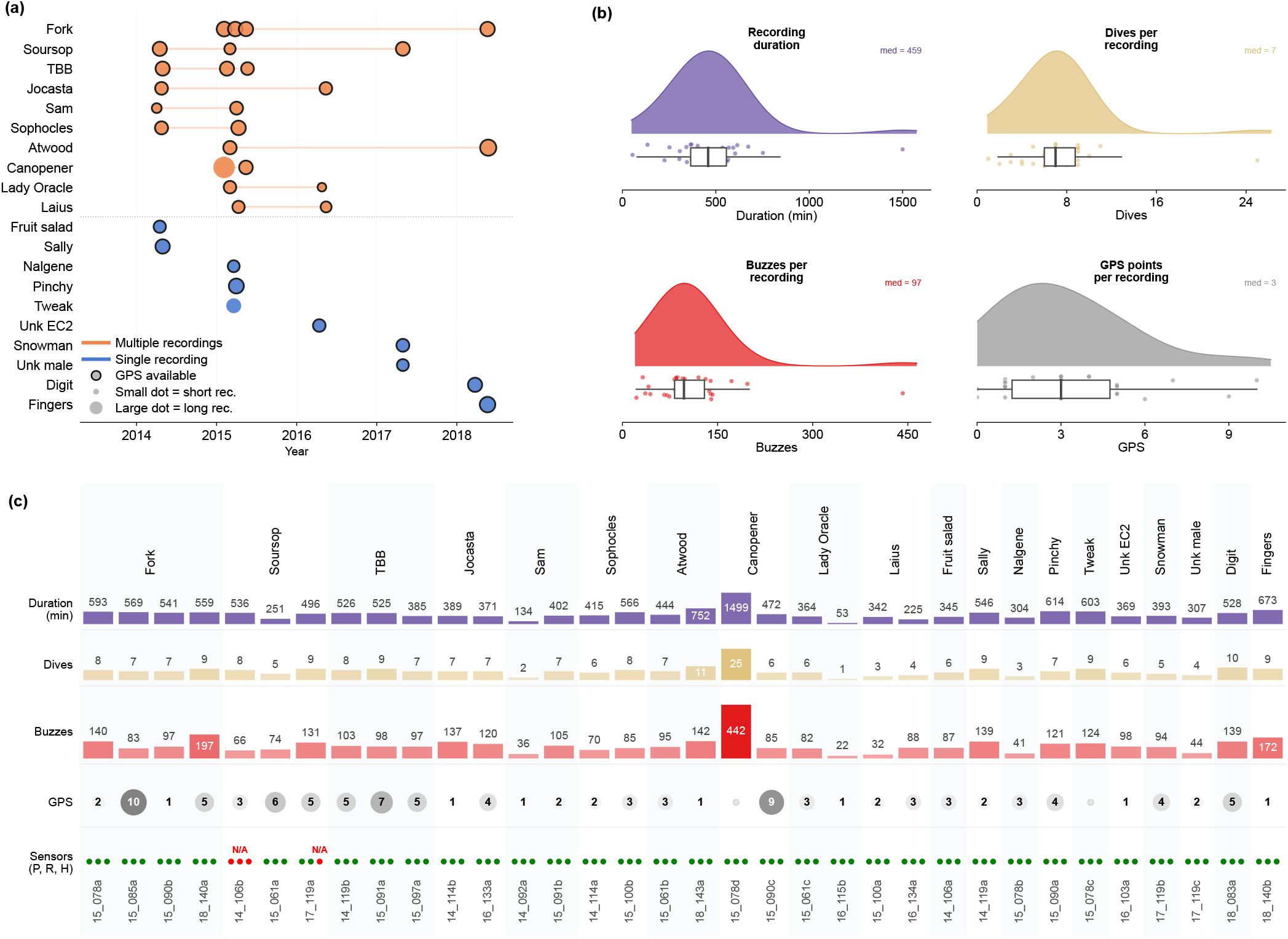
Overview of the DTAG deployments and basic statistics for the Dominica Sperm Whale Project (2014–2018). **(a)** Timeline of tag deployments across individual sperm whales. Orange circles represent individuals tagged multiple times, whereas blue circles denote individuals with a single deployment. Deployments with vessel GPS positions collected while following the surfaced whale are indicated by a dark outline. **(b)** Violin plots showing the distributions of tag duration, number of dives, number of buzzes, and number of vessel GPS positions across deployments. **(c)** Per-deployment summary grouped by individual whale. Horizontal bars show recording duration, number of dives, and number of buzzes. The GPS row gives the number of vessel positions available for each deployment. The bottom row indicates availability of accelerometer- and magnetometer-derived pitch, roll, and heading (P, R, H); red dots indicate sensor failure or unavailable data.

### B. Depth-rate dead-reckoning (Depth-rate DR)

In this method the instantaneous swimming speed *v*(*t*) is derived from the rate of depth change and the instantaneous pitch angle:

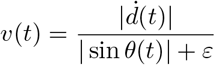

where 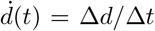 is the numerical depth derivative, and *ε* = 10^−3^ is a small constant that prevents division by zero during near-horizontal swimming segments (|*θ*| ≈ 0). The velocity vector in the global frame is obtained by projecting *v*(*t*) along the forward body axis via the full rotation matrix:

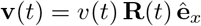

This method approximates the approach used in early DTAG studies [3, 7] and serves as the baseline for comparison. Its principal limitation is that speed estimates become unreliable when the animal swims near-horizontally (e.g., at the surface), leading to large horizontal excursions when pitch is noisy or near zero.

### C. Constant-speed dead-reckoning (Constant-speed DR)

Speed is assumed constant throughout the deployment:

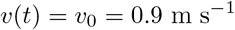

This value is consistent with published mean swimming speeds for sperm whales during foraging dives and on surface [3, 52]. The velocity vector is:

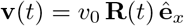

Although this approach ignores within-dive speed variation, it avoids the instability of depth-rate estimation at low pitch. It provides a simple, reproducible benchmark against which more complex methods can be judged.

### D. Pitch-adaptive dead-reckoning (Pitch-adaptive DR)

Prior to integration, depth, pitch, and heading signals are smoothed with a 5-sample moving-average filter to reduce high-frequency sensor noise. This method switches speed estimation strategy per sample based on two simultaneous conditions:

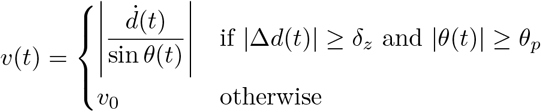

where *δ*_*z*_ = 0.05 m sample^−1^is the depth-change threshold and *θ*_*p*_ = 0.05 rad (≈ 3°) is a secondary pitch safety gate. The dual condition is necessary: a large Δ*d* can still occur with near-zero pitch due to sensor noise, in which case |sin *θ*| would amplify the speed estimate to implausible values. When the whale is actively diving or ascending and pitch is reliably non-zero, speed is derived from vertical kinematics; otherwise a constant fall-back speed *v*_0_ = 0.9 m s^−1^is used. Depth-derived speed estimates are clipped to [0.1, 5.0] m s^−1^to remove physically implausible values. Horizontal position is integrated as

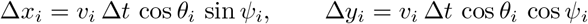

and depth *z* is set directly from the smoothed depth signal, eliminating depth integration drift.

### E. Observed Climb/Descent Rate dead-reckoning (OCDR)

As in pitch-adaptive DR, all sensor signals are pre-smoothed with a 5-sample moving-average filter. The OCDR method (following Miller et al. 2004 and Wensveen et al. 2015) estimates speed when the whale is actively diving or ascending and pitch is sufficiently steep:

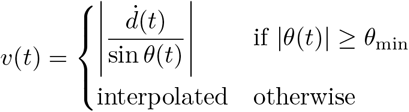

where *θ*_min_ = 0.4 rad (≈ 23°) is the minimum pitch angle below which the OCDR estimate is unreliable. Note that the *ε* regularization used in depth-rate DR is not required here: the threshold *θ*_min_ guarantees |sin *θ*| ≥ sin(0.4) ≈ 0.39 whenever OCDR is applied, keeping the denominator safely away from zero. For near-horizontal segments where |*θ*| *< θ*_min_, the instantaneous speed is estimated by linear interpolation between the nearest preceding and following valid OCDR-derived speed values. If no valid OCDR estimates exist in a recording, the constant fall-back speed *v*_fb_ = 0.9 m s^−1^ is used throughout. Estimated speeds are clipped to [0.1, 5.0] m s^−1^. Horizontal position is integrated with the same projection as pitch-adaptive DR, and depth is again set directly from the smoothed depth signal.

### F. Trajectory integration

For all four methods, the 3D position p(*t*) is obtained by Euler integration:

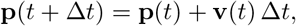

with Δ*t* = 1*/f*_*s*_ = 0.04 s. The initial horizontal position is set to (0, 0), and the initial depth is set to the first recorded depth value after discarding the first 10 samples (to remove sensor equilibration artefacts). A rotation *α* around the vertical axis was applied to each trajectory to correct for the initial whale’s swimming direction. The rotation angle *α* was determined empirically for each recording by a greedy search minimizing the mean nearest-neighbor distance between the reconstructed trajectory and the available vessel GPS points. Rotation angles ranged from 0^◦^to 200^◦^ across deployments.

**Figure S2.**
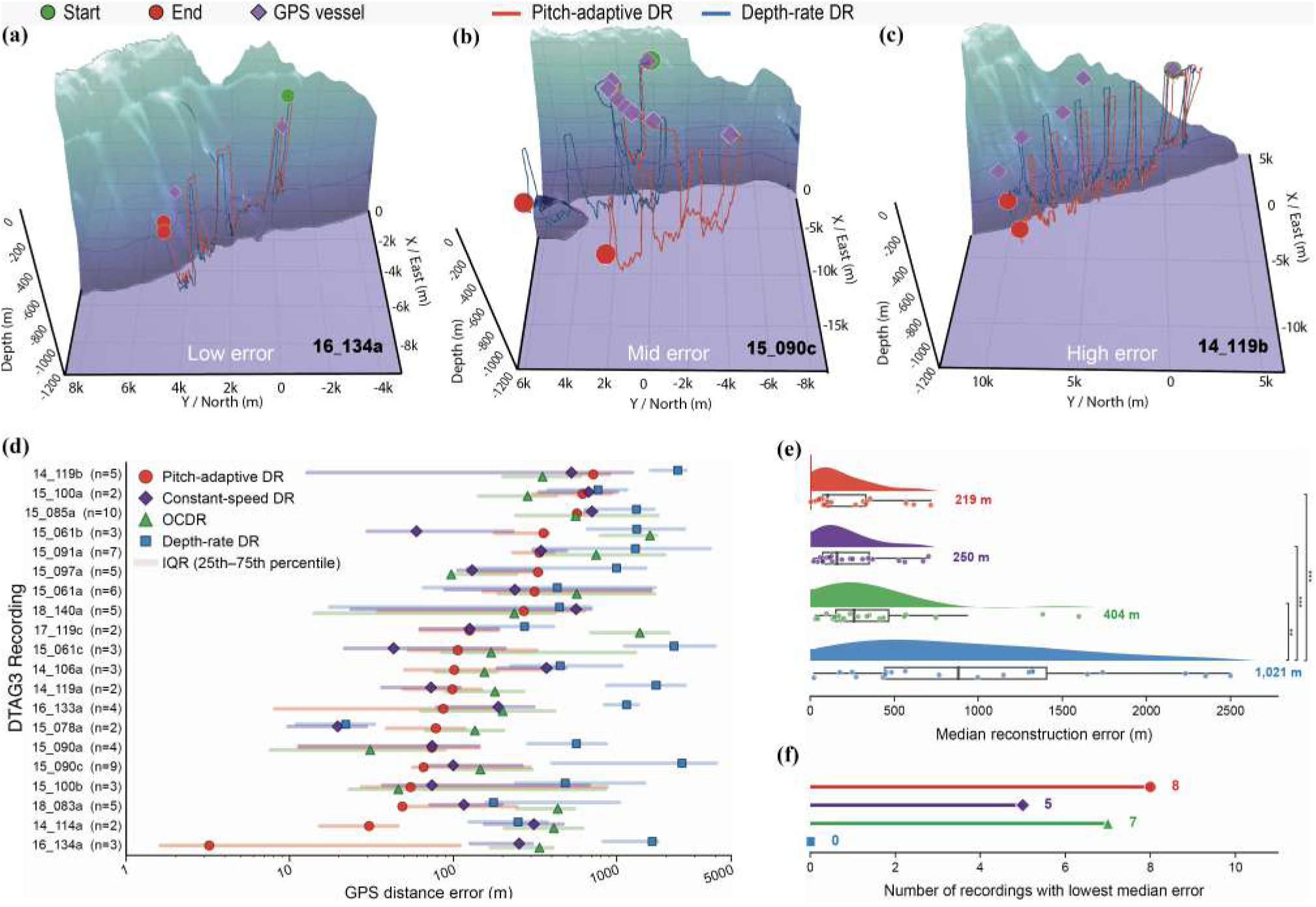
Comparative reconstruction error analysis among dead-reckoning (DR) methods. **(a–c)** Representative 3D reconstructions of whale trajectories for recordings with low (**a**, `16_134a`), moderate (**b**, 15_090c), and high (**c**, `14_119b`) reconstruction errors. Red tracks denote the pitch-adaptive method, while blue tracks denote the depth-rate model, shown relative to surface vessel GPS positions (purple diamonds) used for error assessment. **(d)** Reconstruction error of the four DR models across *N* = 20 DTAG recordings. The value *n* adjacent to each recording ID indicates the number of available GPS positions used for error calculation. Horizontal bars represent the interquartile range (IQR) of GPS distance errors. **(e)** Distribution of median reconstruction errors for all methods. Mann–Whitney *U* tests indicate that the depth-rate DR model (blue) had significantly higher error than all other models (vs. OCDR, *P* = 0.003; vs. constant-speed, *P <* 0.001; vs. pitch-adaptive, *P <* 0.001). No significant difference was observed between the other methods (all *P >* 0.05). **(f)** Frequency of model performance across deployments. The pitch-adaptive model had the lowest mean deployment-level error; pitch-adaptive and constant-speed reconstructions each yielded the lowest mean error in eight deployments. The mean of the deployment-specific median errors for the pitch-adaptive model was 219 m.

## III. TRAJECTORY-SCALE SEARCH STATISTICS AND SPATIAL RECURRENCE

### A. Lévy-like step lengths and superdiffusive movement

To characterize trajectory-scale movement, we analysed the pitch-adaptive three-dimensional reconstructions used throughout the paper. The primary step-length and global mean-squared-displacement (MSD) analyses were restricted to the deep foraging segment between descent and ascent (Supplementary Fig. S3). We retained dives reaching at least 600 m and containing at least 50 trajectory samples in the foraging segment, yielding *n* = 209 dives from 20 whales. Longer-lag MSD analyses additionally used the whole dive excluding the surface phase, because the foraging segment alone did not reliably support multi-minute lags.

#### Turning-point detection

Turning points were identified along each foraging trajectory using an angle-based criterion: a point was labelled a turning point when the angle between consecutive displacement vectors exceeded *θ* = *π/*6 (30^*◦*^). Step length was the Euclidean distance between successive turning points.

#### Step-length fitting

Power-law tails were fitted by continuous maximum likelihood. For each trajectory, the exponent *µ* in *P*(*ℓ*) ∝ *ℓ*^−*µ*^ was evaluated at 30 candidate lower cut-offs spanning the 5th–80th percentiles of the observed step lengths, and the cut-off minimizing the Kolmogorov–Smirnov distance was selected. We use these exponents descriptively and refer to values in the conventional range 1 *< µ* ≤ 3 as Lévy-like, without inferring a unique or optimal search rule.

#### Mean squared displacement analysis

As an independent test for superdiffusive movement, we computed the mean squared displacement (MSD) as a function of time lag *τ* for each foraging trajectory:

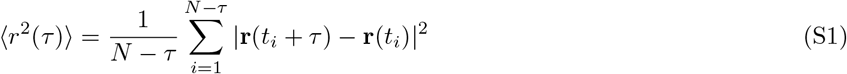

where r(*t*) is the 3D position at time *t* and *N* is the number of trajectory points. Linear regression over logarithmically spaced lags yielded the MSD exponent *α*_MSD_ from 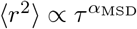. Brownian diffusion has *α*_MSD_ = 1, whereas *α*_MSD_ *>* 1 indicates superdiffusion and *α*_MSD_ = 2 ballistic movement. **Additional results and robustness checks**. The heavy-tailed step-length signature reported in the main text was shared across individuals rather than driven by a few well-sampled whales. Across-whale median *µ* was 2.36, close to the dive-level median of 2.35; most individual distributions overlapped substantially, although estimates from whales represented by few dives remain less precise (Supplementary Fig. S4). The global MSD exponent was similarly consistent across levels of aggregation: median *α*_MSD_ = 1.91 per dive and 1.91 across whales. All dive-level estimates were superdiffusive, and more than 90% exceeded 1.8.

Resolving MSD by timescale revealed a graded crossover rather than a single exponent (Supplementary Fig. S5). In the whole dive excluding the surface phase, movement was nearly ballistic below 1 min (*α* ≈ 1.98), less persistent at 1–5 min (*α* ≈ 1.75), and remained superdiffusive at 5–15 min (*α* ≈ 1.52). At lags of 15 min or more, the estimate remained superdiffusive (*α* ≈ 1.78) but was based on only eight sufficiently long dives and is therefore indicative. The foraging-only trajectory slice agreed at short and intermediate lags but contained too few windows for reliable long-lag inference. Thus, the global *α* ≈ 1.91 summarizes near-ballistic local motion interrupted by turns and less persistent movement over minutes.

This scale dependence was not driven by the treatment of the surface phase. Estimates from the whole dive excluding surface and from the full recording differed by at most 0.03 within supported lag windows (Supplementary Fig. S5). Surface-only movement was near-ballistic below 1 min (*α* ≈ 1.99), and this result was stable as the surface-depth cut-off varied from 5 to 100 m (Supplementary Fig. S6). Including the short surface segment therefore did not generate the dive-scale superdiffusive signature.

Finally, ensemble MSD in a *±*30 s window centred on each classified buzz did not differ between Type A and Type B at any tested lag (paired Wilcoxon signed-rank tests across the eight whales with at least five buzzes of each type; all *P >* 0.05; Supplementary Fig. S7). The A/B contrast is therefore expressed in local kinematics and acoustic timing rather than in the coarser net-displacement field around buzz onset.

Together, the individual-level, timescale, phase-definition, and buzz-centred checks support a robust description of sperm-whale foraging movement as directionally persistent and superdiffusive, with heavy-tailed step lengths consistent with Lévy-like organization. These statistics describe the observed movement structure; they do not establish that whales implement a unique or optimal Lévy search rule, because similar signatures can arise from mixtures of behavioural states, temporal autocorrelation, and environmental heterogeneity [27, 28].

**Figure S3.**
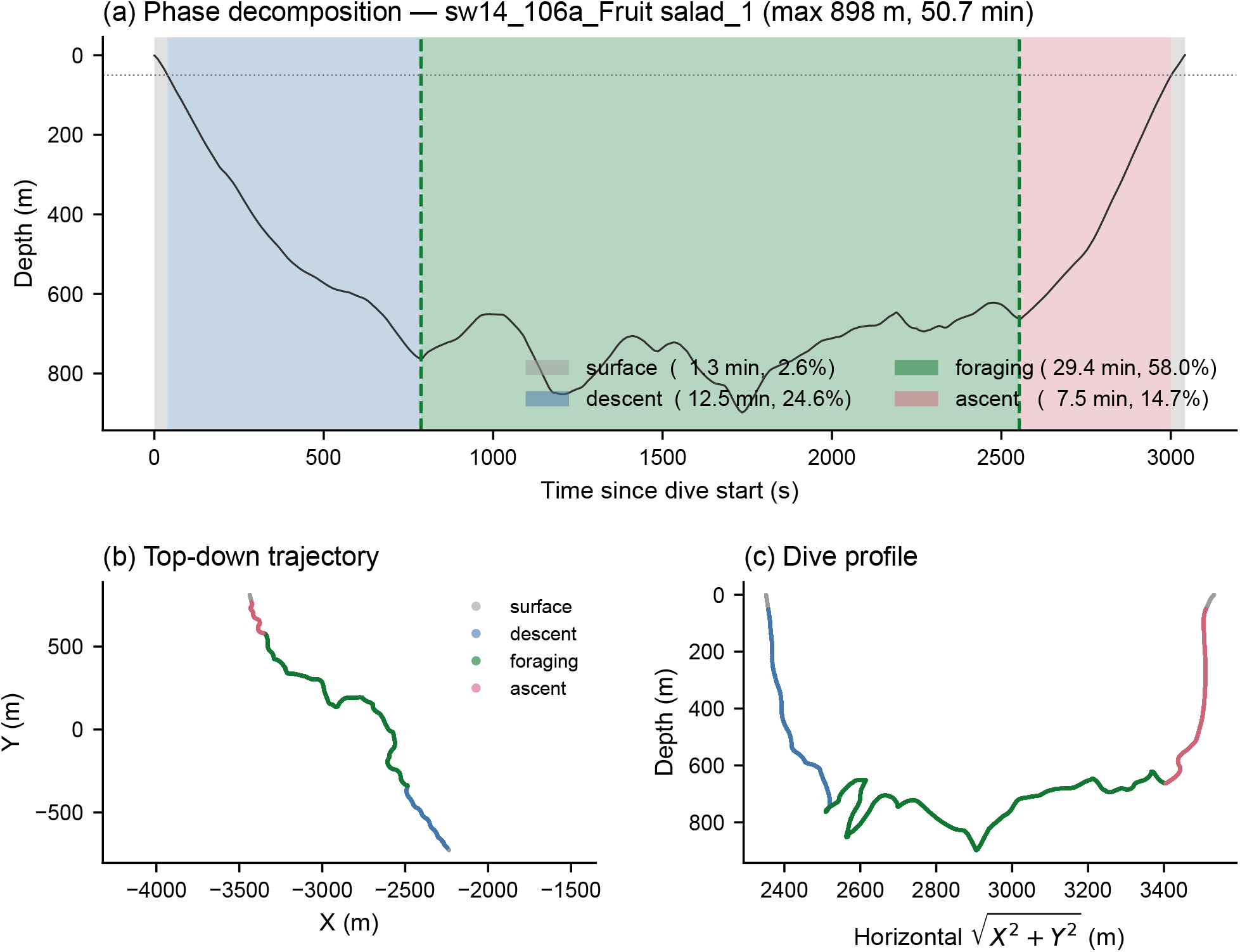
Decomposition of a dive into behavioural periods. Representative deep dive partitioned into surface (depth *<* 50 m), descent, foraging, and ascent. **(a)** Depth profile with dashed lines marking the foraging-window boundaries. **(b)** Top-down trajectory. **(c)** Dive profile against horizontal displacement. Step-length and global-MSD analyses use the foraging period; the alternative trajectory slices used for the timescale robustness analysis are defined in Supplementary Fig. S5.

**Figure S4.**
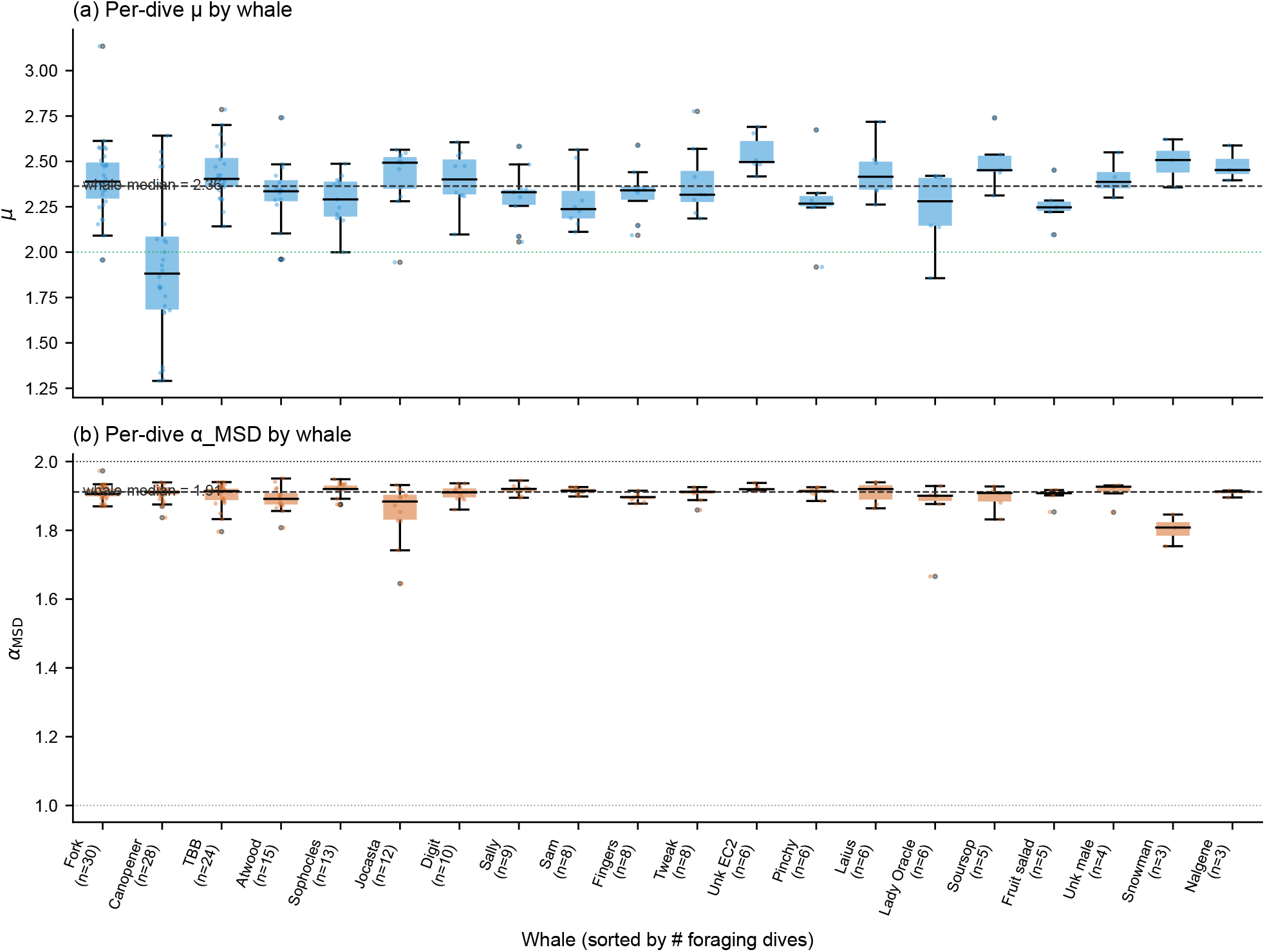
Individual consistency of trajectory-scale statistics. **(a)** Per-dive step-length exponent *µ* for each of the *N* = 20 whales; boxes show the within-whale distribution and points individual dives. The dashed line marks the across-whale median (*µ* = 2.36), and the dotted line marks the reference value *µ* = 2. **(b)** Corresponding global MSD exponents, with an across-whale median of *α*_MSD_ = 1.91. Whales are ordered by the number of retained foraging dives, shown below each name.

**Figure S5.**
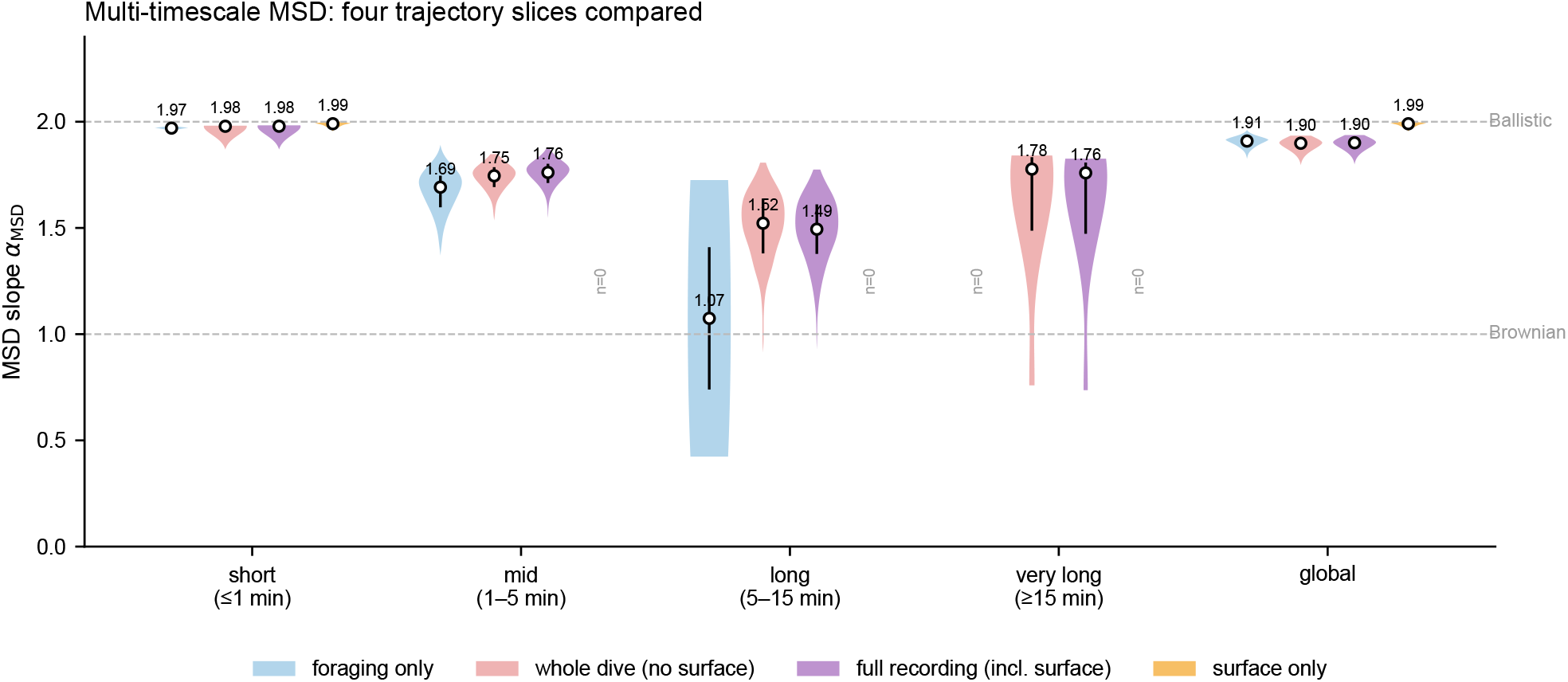
Scale-dependent MSD across alternative trajectory slices. MSD exponent *α*_MSD_ fitted at short (≤ 1 min), intermediate (1–5 min), long (5–15 min), very-long (≥ 15 min), and global scales for the foraging segment, whole dive excluding surface, full recording, and surface-only segment. Circles show medians and vertical lines interquartile ranges. The whole-dive estimate decreases from 1.98 at short lags to 1.75 and 1.52 at intermediate and long lags. Foraging-only trajectories do not provide enough windows for reliable very-long estimates and support only two long-lag fits; those values are shown for completeness. Whole-dive and full-recording estimates are closely aligned, showing that inclusion of surface periods does not generate the scale dependence.

**Figure S6.**
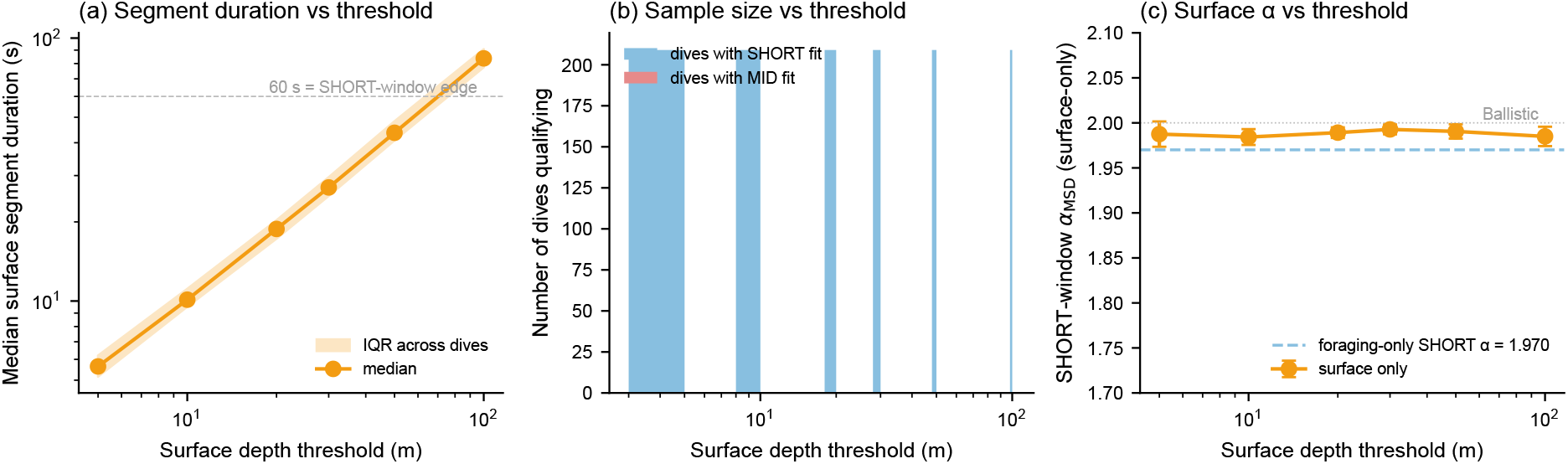
Robustness of the surface characterization to the depth cut-off. **(a)** Median duration of the contiguous surface segment as the depth threshold varies from 5 to 100 m. **(b)** Number of dives supporting short- and intermediate-lag fits. **(c)** Short-window (≤ 1 min) MSD exponent of surface-only motion. Surface movement remains near-ballistic (*α*_MSD_ ≈ 1.99) across all thresholds, showing that the surface result does not depend on the 50-m cut-off used in the primary decomposition.

**Figure S7.**
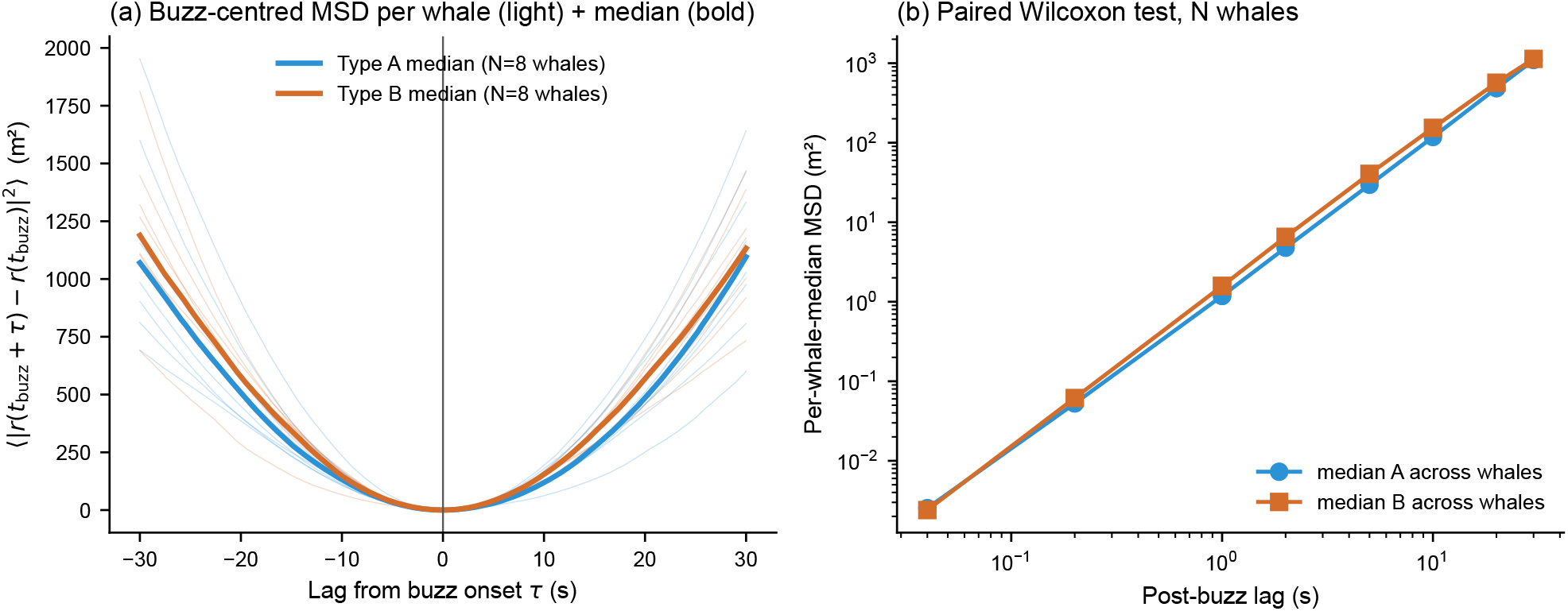
Buzz-centered mean squared displacement does not distinguish prey-capture tactics. Ensemble MSD in a *±*30 s window centered on buzz onset, computed separately for Type A and Type B buzzes and aggregated per individual. **(a)** Per-whale median curves (Type A blue, Type B orange); **(b)** per-lag comparison with per-whale paired Wilcoxon signed-rank tests. No lag shows a significant A-vs-B difference (all *P >* 0.05, *N* = 8 whales with ≥ 5 buzzes of each type), indicating that the Type A/Type B distinction is not expressed in the coarse displacement field around the buzz but in local kinematics and acoustic structure.

**Figure S8.**
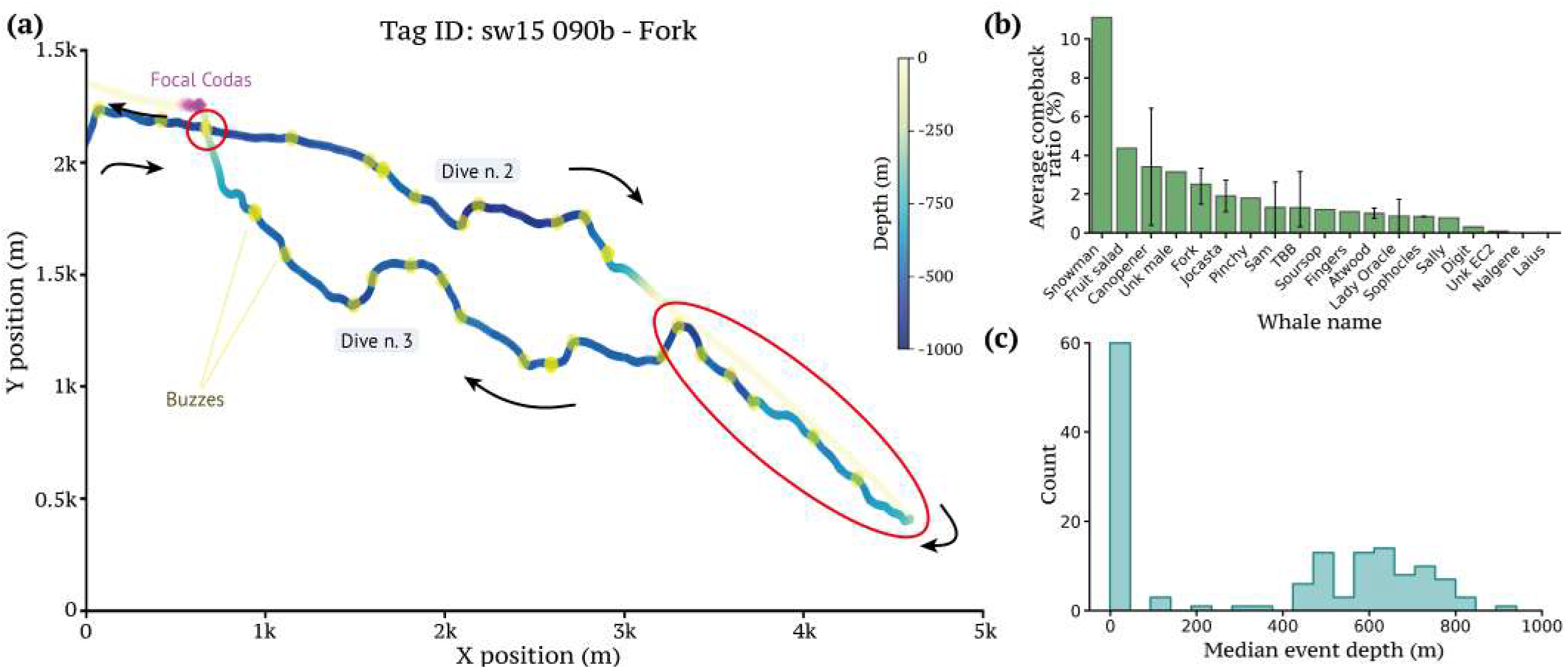
Trajectory with comeback point. **a**. Two-dimensional horizontal trajectory (X vs. Y position) of a part of the recording of the tag `sw15_090b`, for the whale *Fork*. The track line is colored according to depth, ranging from the surface (yellow) to deep waters (dark blue). Black arrows along the path indicate the direction of travel. The trajectory encompasses two distinct dives, which are respectively the second and third dive of the considered recording. Key behavioral events are annotated along the track: purple patches near the end of the trajectory indicate focal codas; yellow dots located along the deeper slopes of the dives denote buzzes, indicative of foraging attempts. Red circles mark the two visits to the same location, and a red star highlights the resulting *comeback point*, defined as a place in the XY plane that the whale visits twice within two dives. **b**. Comeback ratio (proportion of comeback points) across different whales. If one whale has more than one recording, the average and standard deviation across recordings is shown. **c**. distribution of the median depth of each comeback event.

**Figure S9.**
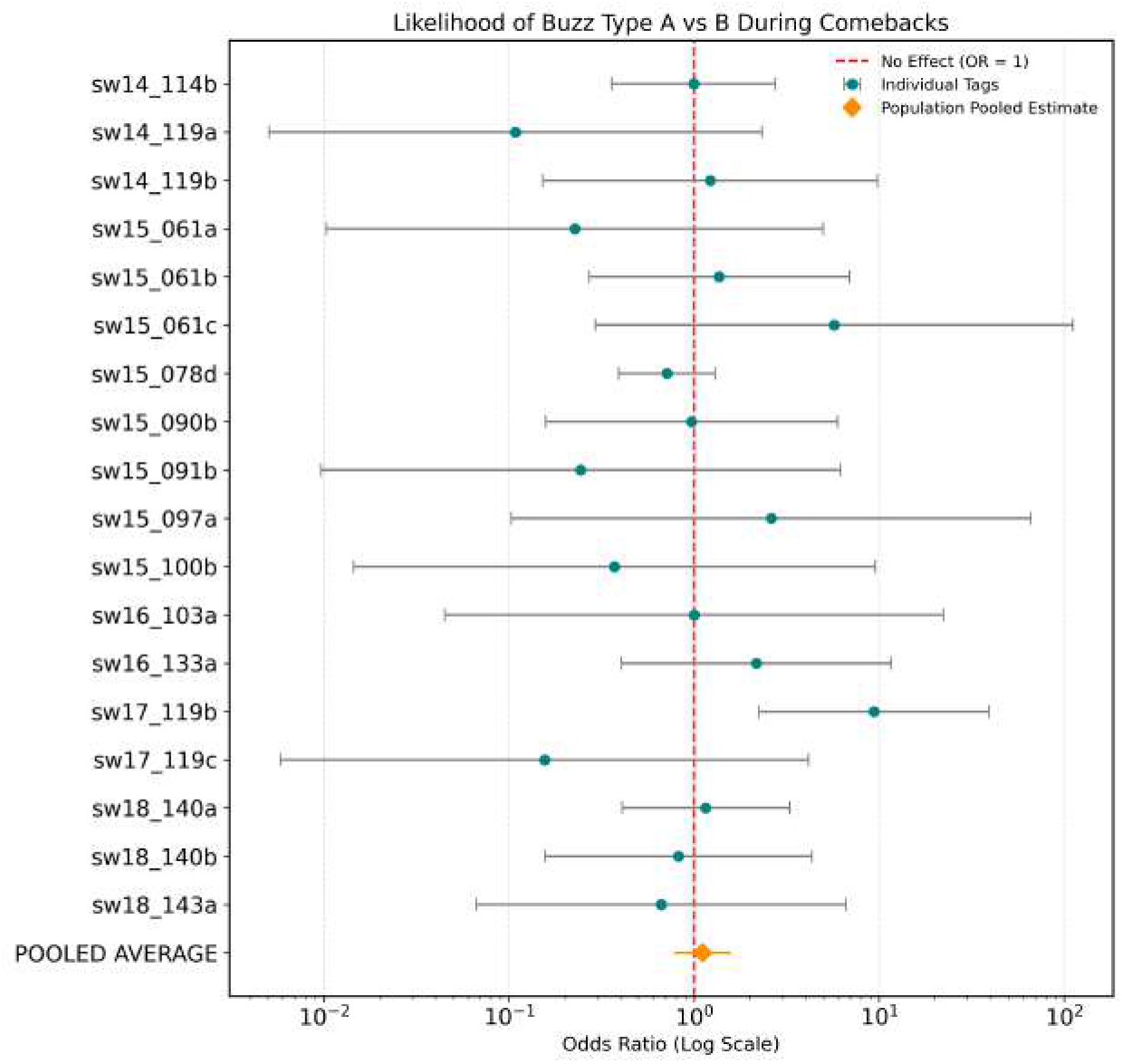
Individual and population-level association between prey-capture tactics and spatial recurrence. Forest plot of the odds of a Type A versus Type B buzz during deep-water comeback loops across 18 deployments. Teal circles show deployment-level odds ratios with 95% confidence intervals; the vertical line marks the null odds ratio of 1. The orange diamond shows the pooled Cochran–Mantel–Haenszel estimate (OR = 1.11, 95% CI 0.78–1.59, *P* = 0.56). The pooled analysis provides no evidence that recurrence is associated with either tactic, although odds ratios were heterogeneous among deployments (*P* = 0.041).

### B. Spatial recurrence and comeback events

We identified spatial recurrence in the horizontal trajectories by pairing each comeback point *P*_*j*_ with the most recent earlier reference point *P*_*i*_ satisfying two conditions: the whale returned within *δ*_close_ = 10 m of the earlier location, and the intervening path had first moved at least *δ*_far_ = 50 m away. This departure criterion excludes local station-keeping and retains genuine returns after a spatial excursion. The comeback ratio for each deployment was defined as the fraction of trajectory samples assigned to such return events and was stable to temporal subsampling up to 60 s.

Comeback ratios ranged from approximately 1% to 30% across deployments. Recurrence occurred both near the surface and at foraging depths of approximately 400–800 m (Fig. S8). To test whether deep-water recurrence was linked to one prey-capture tactic, we compared Type A and Type B buzzes inside and outside comeback intervals, including a 120-s buffer around each interval. A stratified Cochran–Mantel–Haenszel analysis across 18 deployments (*n* = 1,691 classified buzzes) found no population-level association between recurrence and tactic (pooled odds ratio = 1.11, 95% CI: 0.78–1.59, *P* = 0.56; Fig. S9). Spatial return is therefore a feature of broader search organization, but the available data do not tie it to a specific buzz-scale tactic.

## IV. BUZZ CONSTRUCTION, FEATURE SPACE, AND CLASSIFICATION ROBUSTNESS

**Figure S10.**
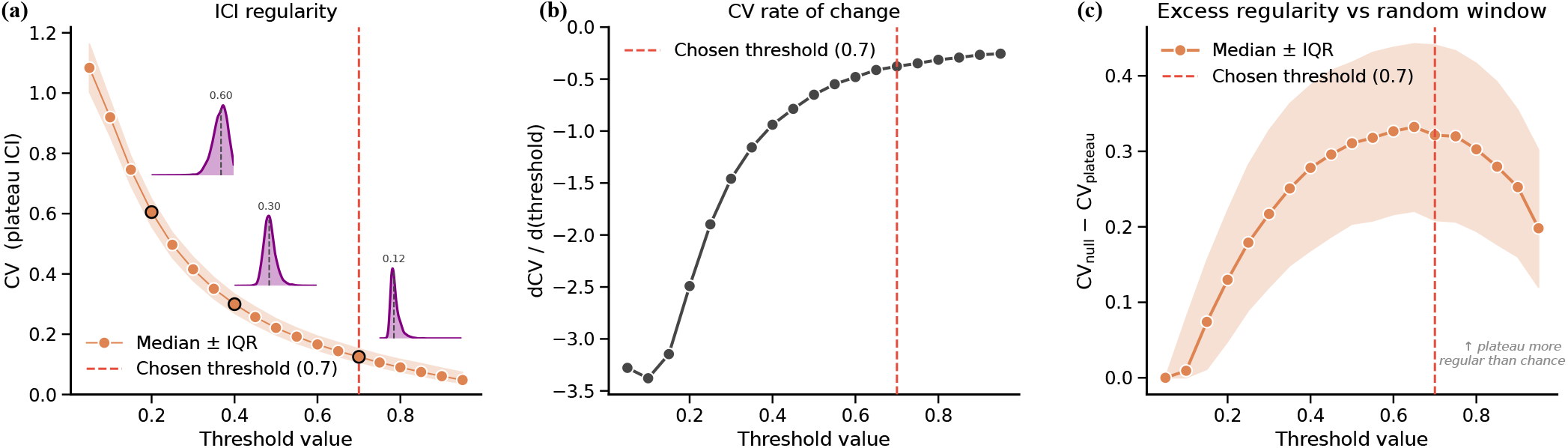
Sensitivity analysis for the choice of plateau detection threshold. Plateau phases within buzzes were detected using a threshold applied to the normalized click-rate range. Two complementary diagnostics were evaluated across thresholds spanning 0.05–0.95, computed over all *N* = 3,686 buzzes. **(a)** Coefficient of variation (CV) of inter-click intervals (ICI) within the detected plateau, shown as median *±* IQR across buzzes. CV decreases monotonically with threshold, reflecting that higher thresholds restrict the plateau to the most rhythmically regular portion of each buzz. Insets show the distribution of per-buzz plateau ICI at thresholds 0.2, 0.4, and 0.7 (median CV: 0.60, 0.30, and 0.12, respectively). **(b)** Numerical derivative of the median CV curve (dCV/d(threshold)). The derivative plateaus near threshold = 0.7, indicating that further increases in threshold yield negligible additional gains in ICI regularity. **(c)** Excess regularity of the detected plateau relative to a null model: for each buzz, 20 same-duration windows were drawn uniformly at random from the full buzz, and the excess is defined as the median null CV minus the plateau CV. Positive values confirm that the detected plateau is genuinely more rhythmically regular than expected by chance. The excess peaks near threshold = 0.7, after which it declines as overly strict thresholds restrict detection to too few clicks. Together, panels (a–c) indicate that a threshold of 0.7 (red dashed line) identifies plateaus that are both internally regular and maximally distinct from the surrounding buzz structure.

**Figure S11.**
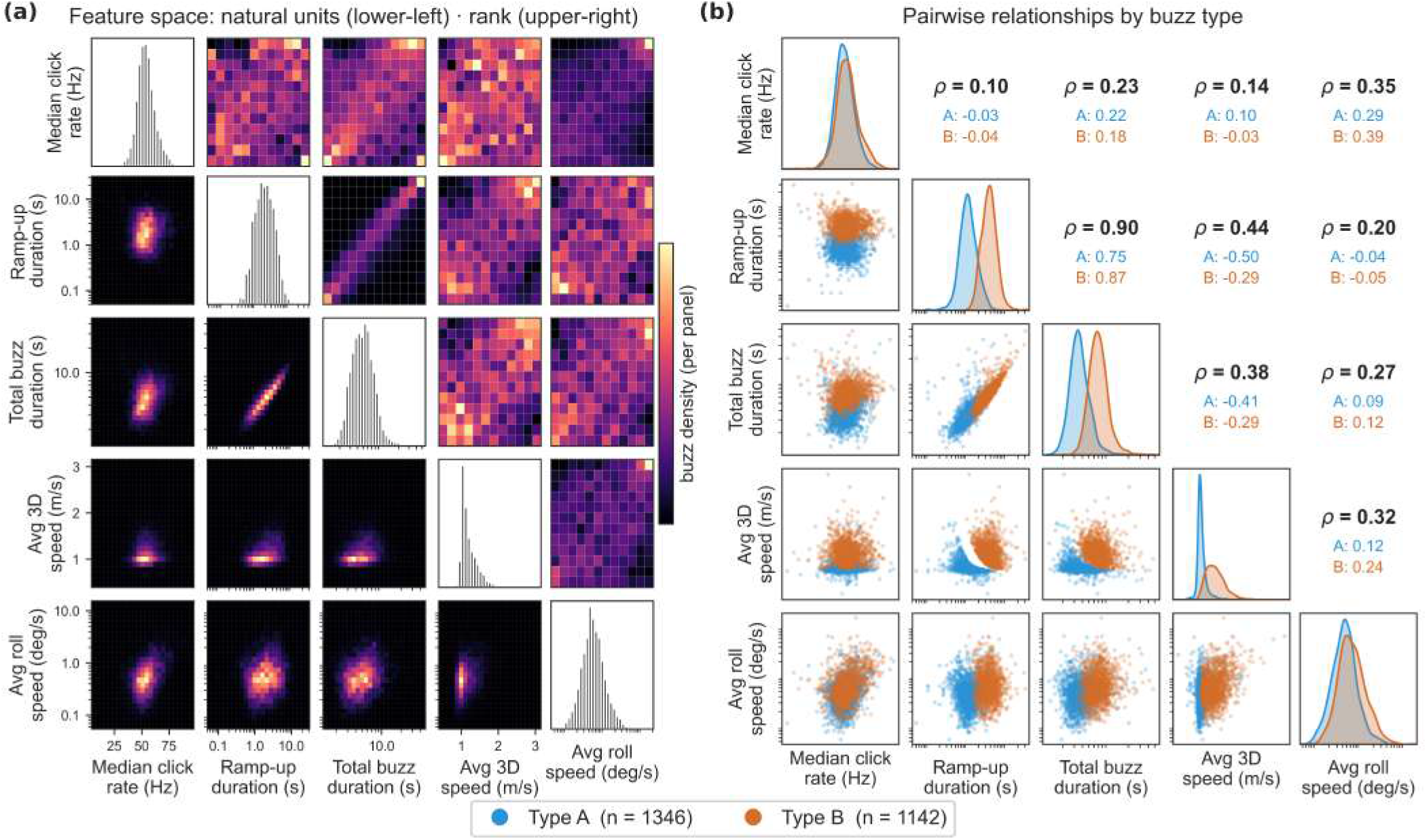
Multivariate structure of the five buzz-characterization features. Each prey-capture buzz is summarized by three acoustic features (median click rate, ramp-up duration, total buzz duration) and two concurrent kinematic features (average 3D swim speed and average roll speed). **(a)** Matrix of pairwise two-dimensional densities for the *N* = 3,488 buzzes with complete feature data (of 3,686 total; 198 lacked a 3D-speed estimate). Each off-diagonal panel is a 2D histogram (brighter = higher buzz density, scaled independently per panel). The lower-left triangle shows the features in natural units (logarithmic axes and tick values for the three right-skewed features: ramp-up duration, total buzz duration and roll speed), while the upper-right triangle shows the same pairs after a quantile (rank) transform that removes the disparate marginal scales and shows the underlying dependence structure; diagonal panels show the one-dimensional marginal distributions. The only strong pairwise association is between ramp-up and total buzz duration (Spearman *ρ* = 0.90, expected as the latter contains the former); all remaining pairs are weakly correlated, indicating the five features carry largely complementary information. **(b)** The same feature pairs resolved by prey-capture tactic, as assigned by a two-component Gaussian mixture model (Type A, blue, *n* = 1,346; Type B, orange, *n* = 1,142; 1,000 ambiguous buzzes below the posterior-probability threshold excluded). Lower-left panels show the bivariate distributions colored by type, diagonal panels show the per-type marginals, and upper-right panels report the Spearman rank correlation for the pooled classified buzzes (black) and within each type (colored). The two tactics separate most clearly along the ramp-up-duration and 3D-swim-speed axes, whereas their median-click-rate and roll-speed marginals largely overlap, confirming that the acoustic–kinematic dichotomy is driven primarily by one acoustic (ramp-up duration) and one kinematic (swim speed) feature.

In the main analysis, we employed a Gaussian mixture model (GMM) approach to classify buzz events into two types based on pairs of acoustic and kinematic features. Prior to clustering, feature pairs were preprocessed using a quantile transform to mitigate the influence of outliers and enforce the Gaussian distribution assumptions required by the model. We fitted a two-component GMM with full covariance matrices, initialized via K-means. We performed the robustness analysis with no threshold on the posterior probability of membership (*p >* 0.), thereby assigning every valid buzz event to its most likely cluster.

As an alternative classification approach, we employed an extreme-corner classification in the same rank-transformed feature plane. Type A is defined as the *N* buzzes with the smallest Euclidean distance from the lower-left corner (0, 0) and Type B as the *N* buzzes closest to the upper-right corner (1, 1).

We performed three pairwise comparisons to evaluate the consistency of buzz-type assignments across methods and feature choices (Supplementary Fig. S12):

1. **Different speed metric, same method** (Supplementary Fig. S12**a**): The GMM fitted with 3D speed was compared with the GMM fitted with vertical speed. Agreement was 95.3% (with Cohen’s *κ* = 0.905) for *n* = 3488 events. Of the 165 discordant events, 95 events classified as Type A by the GMM 3D speed were assigned Type B by the GMM vertical speed, while 70 events showed the opposite pattern. The KDE plots confirm that the two GMM variants divide the feature space similarly, with minor boundary differences in the transition zone.
2. **Same speed metric, different method** (Supplementary Fig. S12**b**): The GMM fitted with 3D speed was compared with the extreme-corner classification using 3D speed using the same feature pair (Duration of ramp-up *×* 3D speed). Agreement was 97.8% (with Cohen’s *κ* = 0.956) for *n* = 3488 events, indicating near-perfect concordance between the two methods. The confusion matrix revealed that all 1787 Type A assignments by the Extreme-corner classification were also classified as Type A by the GMM 3D speed, with only 77 events labeled as Type B by the Extreme-corner classification and Type A by the GMM 3D speed.
3. **Azorean buzz classification’s method vs main analysis’ method** (Supplementary Fig. S12**c**): The GMM fitted with 3D speed was compared with the extreme-corner classification using vertical speed (the feature combination available for the Azorean dataset, which lacks 3D speed measurements). Agreement was 94.5% (with Cohen’s *κ* = 0.890) for *n* = 3488 events. Of the 198 discordant events, 148 events classified as Type A by the GMM 3D speed were assigned Type B by the GMM vertical speed, while 44 events showed the opposite pattern, reflecting a boundary shift when substituting the 3D with the vertical speed. Despite this asymmetry, the overall agreement remained high, supporting the validity of using vertical speed as a proxy when 3D speed data are unavailable.

Across all three comparisons, Cohen’s *κ* exceeded 0.89, confirming that buzz-type classifications are robust to the choice of the clustering algorithm (GMM vs. extreme-corner classification) and speed metric. The KDE plots in Supplementary Fig. S12 further illustrate that the spatial distribution of Type A and Type B events in the rank-transformed feature space is consistent across classification methods.

To further assess the robustness of our buzz-type dichotomy, we applied the extreme-corner classification method to the full DSWP dataset using vertical speed as the kinematic feature (duration of ramp-up rank *×* vertical speed rank, both rescaled to [0, 1]). This configuration mirrors the feature set available for the Azorean dataset, providing a direct test of whether the behavioral patterns identified by the GMM are recoverable by a fundamentally different, geometry-based classification scheme.

Supplementary Fig. S18 presents the full set of behavioral analyses under this alternative classification, reflecting the structure of Figs. 3 and 4 in the main text. The bivariate density of rank-transformed features (Supplementary Fig. S18**a**) preserves the bimodal structure observed under the GMM, with type A events concentrated near the origin (short ramp-up, low vertical speed) and type B events clustered toward the upper-right corner (long ramp-up, high vertical speed). Both buzz types are represented across the majority of tagged individuals (Supplementary Fig. S18**b**), confirming that the behavioral dichotomy is not driven by individual identity.

All key kinematic and acoustic features reported in the main text are reproduced. Type A buzzes are preceded by unimodal, low-intensity vertical speeds, whereas Type B buzzes exhibit the characteristic bimodal distribution indicative of high-speed vertical pursuit (Supplementary Fig. S18**c**). Roll speed is significantly higher for Type B events in the seconds around buzz onset (Supplementary Fig. S18**d**), consistent with rotational prey-capture maneuvers. The proportional distribution of vertical movement classes (Supplementary Fig. S18**e**) preserves the marked shift toward persistent vertical directionality in Type B (76%) compared to the more frequent oscillatory patterns in Type A (43%). The Markov chain transition structure (Supplementary Fig. S18 f) remains qualitatively unchanged, Type B events are approximately twice as likely to terminate a dive sequence (0.09 *±* 0.01) compared to Type A (0.05 *±* 0.01), and the self-transition and cross-transition probabilities are qualitatively consistent with those obtained under the GMM. Pre-buzz path straightness is significantly higher for Type B (Supplementary Fig. S18**g**), while post-buzz straightness shows a negligible effect size (Supplementary Fig. S18**h**). The distribution of inter-buzz pause durations (Supplementary Fig. S18**i**) further confirms a significant association between prey-capture tactic and temporal spacing of attempts.

Together, these results demonstrate that the two prey-capture tactics and their associated kinematic signatures are not artifacts of the GMM clustering procedure, but reflect a robust behavioral structure recoverable across classification methods.

**Figure S12.**
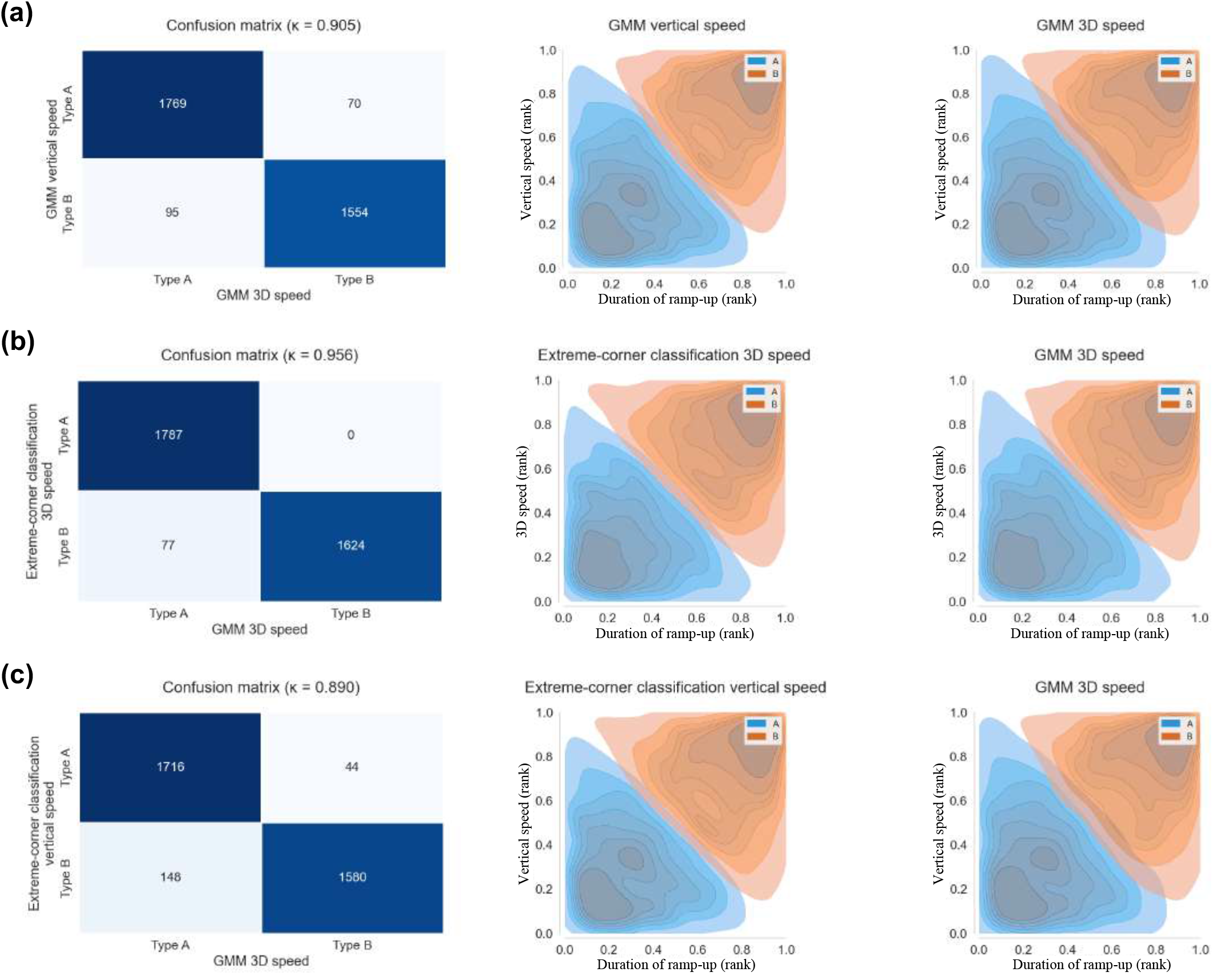
Robustness of buzz clustering across alternative classification schemes. Each row compares an alternative classifier with the primary GMM using 3D swim speed. The left column shows the confusion matrix and Cohen’s *κ*; the centre and right columns show bivariate densities under the alternative and primary classifiers. Density plots use empirical ranks rescaled to [0, 1]. **(a)** Alternative GMM using vertical speed (agreement = 95.3%, *κ* = 0.905). **(b)** Extreme-corner classification using 3D speed (agreement = 97.8%, *κ* = 0.956). **(c)** Extreme-corner classification using vertical speed (agreement = 94.5%, *κ* = 0.890). All comparisons use the *n* = 3488 buzzes with complete feature data; no posterior-probability threshold was applied in this robustness analysis.

**Figure S13.**
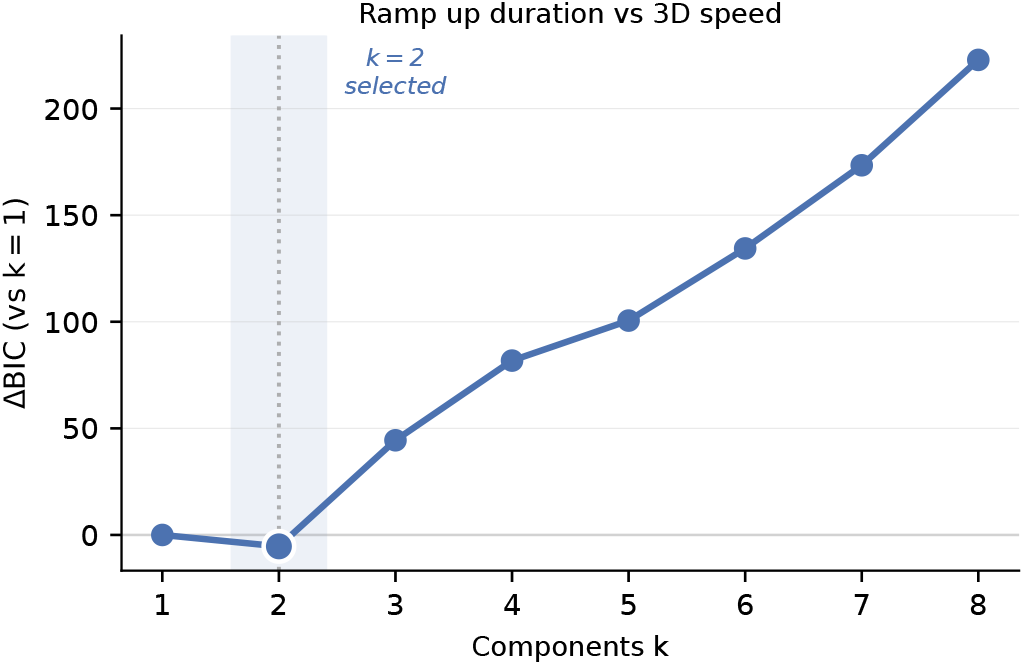
BIC model selection for Gaussian Mixture Model clustering of buzz kinematics. Δ BIC relative to a single-component model (*k* = 1) as a function of the number of GMM components *k*, fitted to the (ramp-up duration, 3D swim speed) feature plane. The minimum BIC is achieved at *k* = 2 (dashed line, shaded region), indicating that two components provide the best trade-off between model fit and complexity. BIC increases monotonically for *k >* 2, confirming that additional components do not improve model fit. The two-component solution (*k* = 2) was therefore selected for all downstream buzz classification analyses described in the main text.

**Figure S14.**
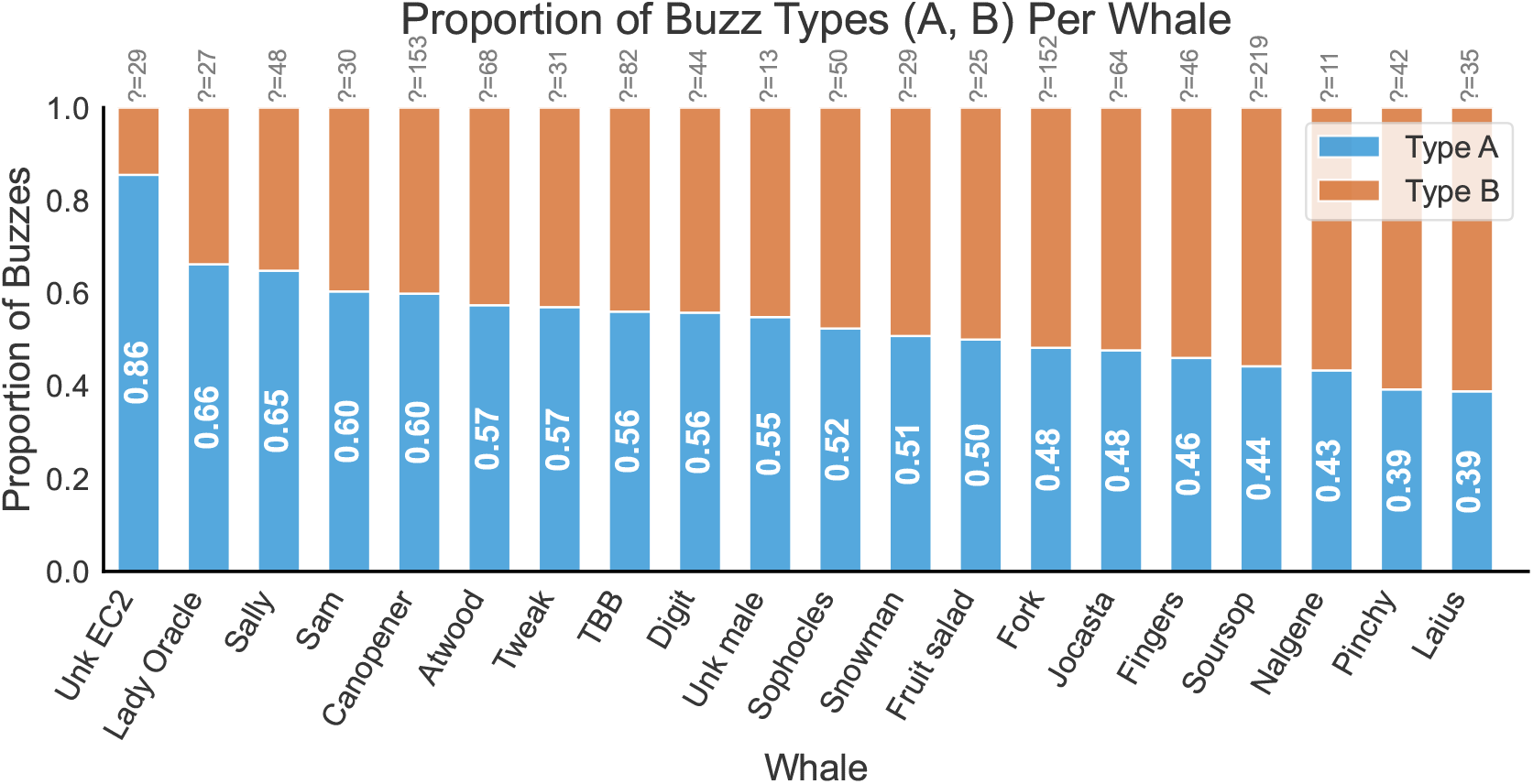
Individual variation in buzz-type composition. Normalized stacked bars show the proportion of Type A (blue) and Type B (orange) buzzes for each individual whale, sorted by decreasing Type A fraction. White numerals indicate the Type A proportion; numbers above each bar denote buzzes excluded because posterior cluster probability was below 0.75. Type A proportions range from 0.86 (Unk EC2) to 0.39 (Pinchy and Laius), with most sampled individuals showing a moderate Type A majority.

**Figure S15.**
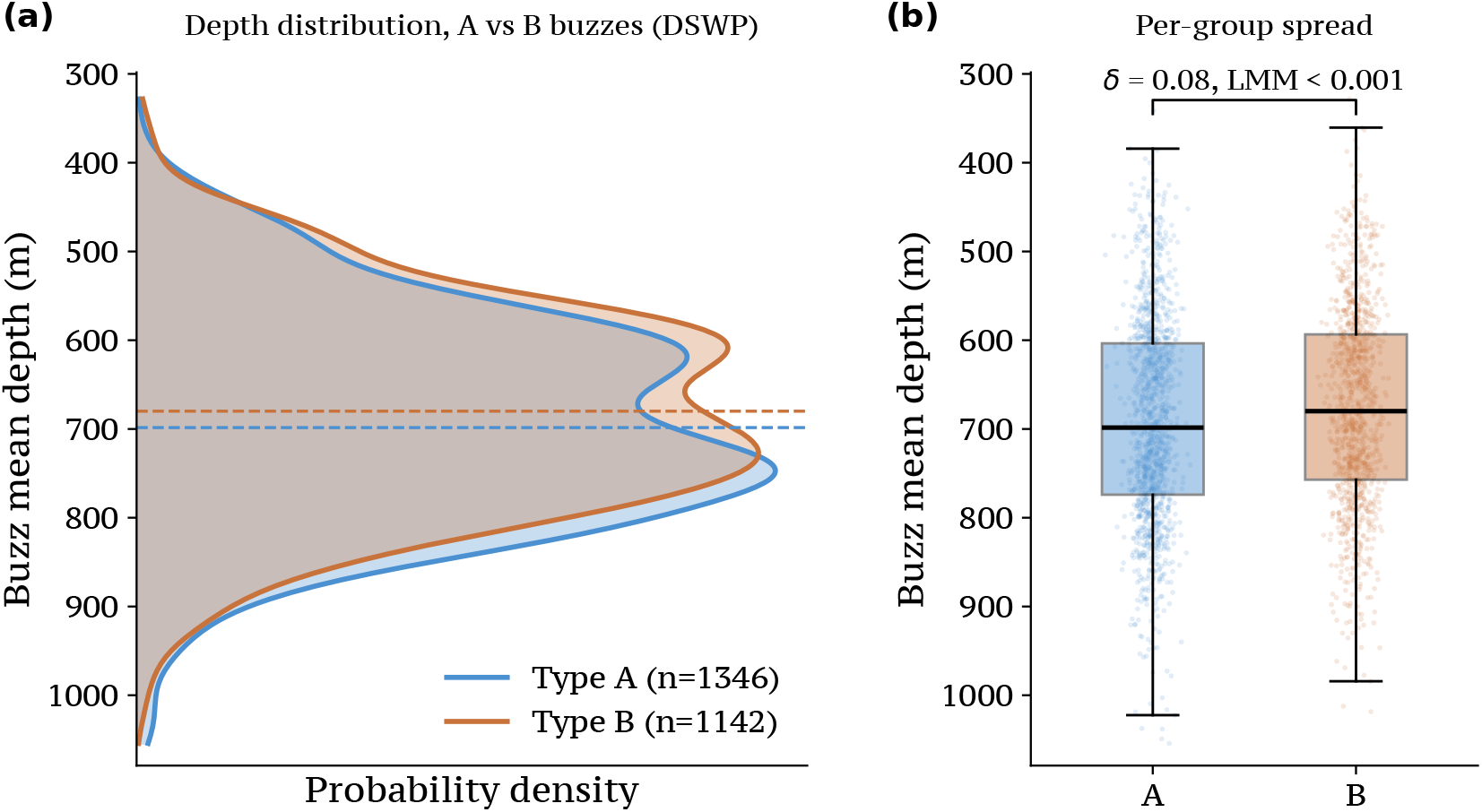
Depth distribution of Type A and Type B buzzes (DSWP). **(a)** Probability density of buzz mean depth for Type A (blue, *n* = 1346) and Type B (orange, *n* = 1142) buzzes; dashed lines indicate group medians (698.5 and 679.9 m). The two distributions overlap across the full ~ 330–1050 m foraging range. **(b)** Per-group spread of buzz mean depth (boxes: median and interquartile range; whiskers: 1.5*×*IQR; points: individual buzzes). A linear mixed-effects model with buzz type as a fixed effect and random intercepts for individual whale and for dive nested within whale indicated a statistically significant but negligible difference (estimate +10.6 m, 95% CI 4.5–16.8 m; LMM *p <* 0.001; Cliff’s *δ* = 0.08). The depth axis is inverted so that greater depths appear lower. Together, the panels show that depth does not distinguish the two buzz tactics.

## V. ENERGETIC, GEOMETRIC, AND SEQUENCE-LEVEL CHARACTERIZATION

### A. Energy-expenditure model

#### Source and conceptual framework

The per-buzz values reported in Fig. 3**g** were computed with a trajectory-driven time-series model inspired by the steady-swimming formulation of Aoki et al. [30] and the marine-mammal raptorial-capture framework of Goldbogen et al. [31], and modified to account for motor demand using the reconstructed trajectories. The energetic model combines work against parasite drag, work performed during positive tangential acceleration, and baseline rest-of-body metabolism over the observed buzz. The analytical Goldbogen model requires event-specific prey speed, reaction distance, interception timing, and capture geometry, none of which were measured in this dataset; we therefore use its hydrodynamic and energetic framework without claiming a literal fit of its prey-interception model.

#### Governing equations and application

For each buzz, speed *V* (*t*) was calculated from successive positions in the pitch-adaptive 25-Hz reconstructed trajectory over the full interval from the first to the last buzz click. Both acoustic boundaries were mapped to the preceding trajectory sample, and tangential acceleration was calculated as *a*(*t*) = *dV* (*t*)*/dt*. Parasite drag was

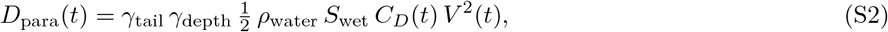

where *ρ*_water_ is seawater density, *S*_wet_ is wetted surface area, and *γ*_tail_ and *γ*_depth_ are tail-heaving and depth correction factors. The instantaneous drag coefficient was

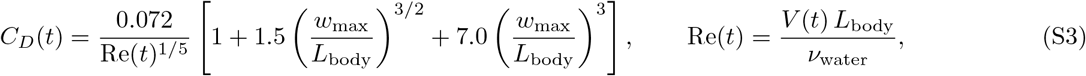

where *L*_body_ is body length, *w*_max_ is maximum skull width, and *ν*_water_ is kinematic viscosity. Wetted surface area is estimated allometrically as 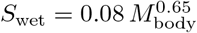.

Work against parasite drag and during positive acceleration was integrated as

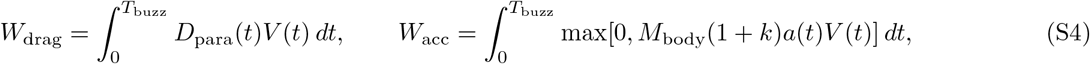

where *k* is the added-mass coefficient. Deceleration was assigned no negative or recovered energetic work. The metabolic equivalent of this mechanical work was

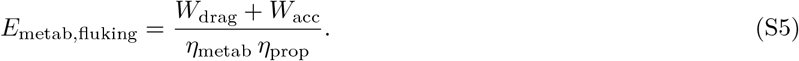

Rest-of-body metabolism was integrated over the same observed interval:

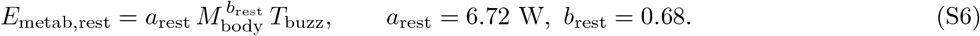

The total per-buzz metabolic energy is the sum, *E*_total_ = *E*_metab,fluking_ + *E*_metab,rest_.

#### Parameter table

The fixed sperm-whale parameters used in the production run are listed in Table S1.

For the comparison shown in Fig. 3**g**, Type A and Type B values were first averaged within each whale, pooling repeated deployments, and the resulting paired individual means were compared across *N* = 20 whales using a paired t-test.

**Table S1.** Parameters used in the trajectory-driven energetic model, inspired by the formulations of Aoki et al. [30] and Goldbogen et al. [31]. The same fixed morphology was used for Type A and Type B buzzes.

| Parameter | Symbol | Value (units) | Source / notes |
| --- | --- | --- | --- |
| Body length | $L_{\text{body}}$ | 12.0 m | fixed population-level value |
| Body mass | $M_{\text{body}}$ | $4.0 \times 10^4$ kg | fixed population-level value |
| Max skull width | $w_{\text{max}}$ | 1.5 m | fixed population-level value |
| Wetted surface area | $S_{\text{wet}}$ | $0.08 M_{\text{body}}^{0.65}$ | allometry, [31] |
| Seawater density | $\rho_{\text{water}}$ | $1027 \text{ kg m}^{-3}$ | standard |
| Kinematic viscosity | $\nu_{\text{water}}$ | $1.15 \times 10^{-6} \text{ m}^2 \text{ s}^{-1}$ | seawater $\approx 10^\circ\text{C}$ |
| Metabolic efficiency | $\eta_{\text{metab}}$ | 0.25 | mammalian standard [31] |
| Propeller efficiency | $\eta_{\text{prop}}$ | 0.75 | cetacean range 0.7–0.8 |
| Tail-heaving correction | $\gamma_{\text{tail}}$ | 2.5 | range 2–3 |
| Depth correction | $\gamma_{\text{depth}}$ | 1.0 | deep foraging (not surface) |
| Added-mass coefficient | $k$ | 0.06 | effective accelerating mass |
| Resting metabolic constant | $a_{\text{rest}}$ | 6.72 W | allometric |
| Resting metabolic exponent | $b_{\text{rest}}$ | 0.68 | mass-scaling exponent |

#### Alternative model formulations

We evaluated the sensitivity of the A–B contrast using the steady-drag formulation of Aoki et al. [30], which estimates the metabolic equivalent of active-swimming drag as

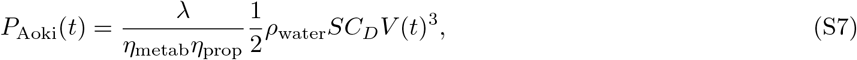

using an active-to-passive drag ratio *λ* = 3 and a constant drag coefficient *C*_*D*_ = 0.00306. We integrated this expression over the same reconstructed speed series and first-click–last-click intervals used by the energetic model. To isolate the effect of model formulation, the primary comparison used the same allometric surface area and conversion efficiencies as Table S1. Across individual means, the Aoki formulation yielded 0.013 MJ for Type A and 0.111 MJ for Type B (*B/A* = 8.51; paired t-test, *P* = 5.1 *×* 10^−13^), with Type B higher in all 20 whales. Using Aoki et al.’s original surface area of 23.7 m^2^rescales the corresponding values to approximately 0.004 and 0.034 MJ but leaves the ratio unchanged. Thus, removing the positive-acceleration term changes the absolute values but preserves both the direction and approximate scale of the energetic contrast.

We did not include a numerical comparison with the analytical prey-interception model of Goldbogen et al. [31]. Fitting that model requires event-specific prey speed, reaction distance, interception timing, and capture geometry, which are unavailable here. Assigning fixed values or substituting whale onset speed would not constitute a fitted application of the model and could impose or obscure a tactic difference through unobserved inputs. We therefore retain Goldbogen et al. as a source for the hydrodynamic framework, but not as a separate quantitative comparison.

**Table S2.** Comparison of energetic formulations. Values are means of individual Type A and Type B means. Although the two formulations assign different absolute values, both recover a marked energetic asymmetry between tactics, with Type B higher in every whale and a similar B/A ratio.

| Formulation | Type A (MJ) | Type B (MJ) | B/A | Quantity represented |
| --- | --- | --- | --- | --- |
| Energetic model | 1.275 | 9.069 | 7.11 | Drag + positive acceleration + rest over observed buzz |
| Aoki steady-drag model | 0.013 | 0.111 | 8.51 | Active-swimming drag over observed buzz |

#### Component contributions, limitations, and interpretation

**Figure S16.**
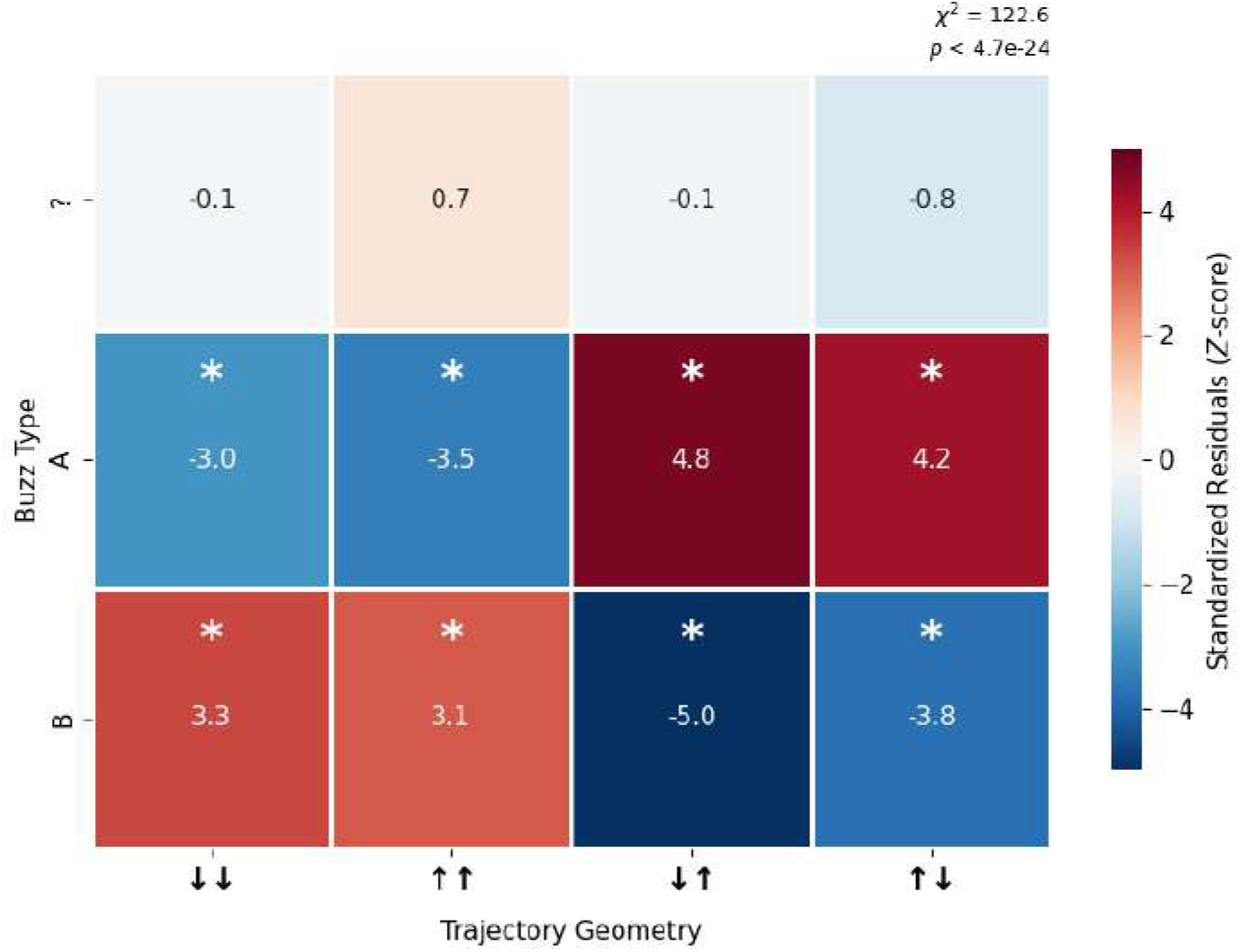
Standardized residuals evaluating the association between acoustic Buzz Type and Trajectory Geometry. A chi-square test of independence revealed a highly significant relationship between the variables (*χ*^2^ = 122.6, *P <* .001). Cell colors represent standardized residuals (*Z*-scores), indicating the direction and magnitude of the deviation from expected frequencies. Red cells represent a higher-than-expected occurrence, while blue cells represent a lower-than-expected occurrence. Asterisks (*) denote specific buzz-geometry combinations that significantly deviate from expected values (|*Z*| ≥ 1.96, *P <* .05). We see that buzzes of type A are predominantly associated with oscillatory maneuvers, whereas B buzzes are strongly associated with persistent vertical directionality.

In the energetic model, the mean metabolic equivalents of drag, positive acceleration, and resting metabolism were respectively 0.010, 1.229, and 0.037 MJ for Type A, and 0.076, 8.924, and 0.070 MJ for Type B. Positive acceleration consequently contributes 96.3% and 98.4% of the respective total estimates, as expected for a model designed to represent motor demand during the observed buzz trajectory. Differentiation of a 25-Hz dead-reckoned speed series can amplify high-frequency reconstruction noise, and retaining positive work while assigning no recovered work to deceleration prevents positive and negative fluctuations from cancelling. For this reason, the estimates are most informative as a relative comparison between tactics. The Aoki formulation, which omits the acceleration term entirely, yields a similar B/A ratio (8.51 rather than 7.11) and preserves Type B above Type A in every whale. The marked energetic separation is therefore not generated solely by the treatment of positive acceleration. Three further caveats deserve explicit treatment:

1. **Fixed morphology**. The model uses a common body length and mass for all buzzes (Table S1), whereas the DSWP cohort is predominantly female and immature. Because Type A and Type B use identical morphology within each whale, the paired A/B contrast and within-whale rank order are less sensitive to this assumption than the absolute energy values, which should be interpreted as model-dependent estimates.
2. **Incomplete mechanical budget**. The model represents translational drag and acceleration but not rotational work, turning, buoyancy, lift, internal fluking work, or variation in propulsive efficiency. It therefore captures only part of the energetic budget for prey capture.
3. **Event boundaries and prey inputs**. The first and last buzz clicks define an observable, reproducible interval but may not delimit the complete pursuit. Prey speed, reaction distance, capture success, and prey energetic content were not measured and are not inferred from the energetic estimates.

The complete implementation of the energetic model and the Aoki comparison will be included in the public repository described in the Data and code availability statement.

**Figure S17.**
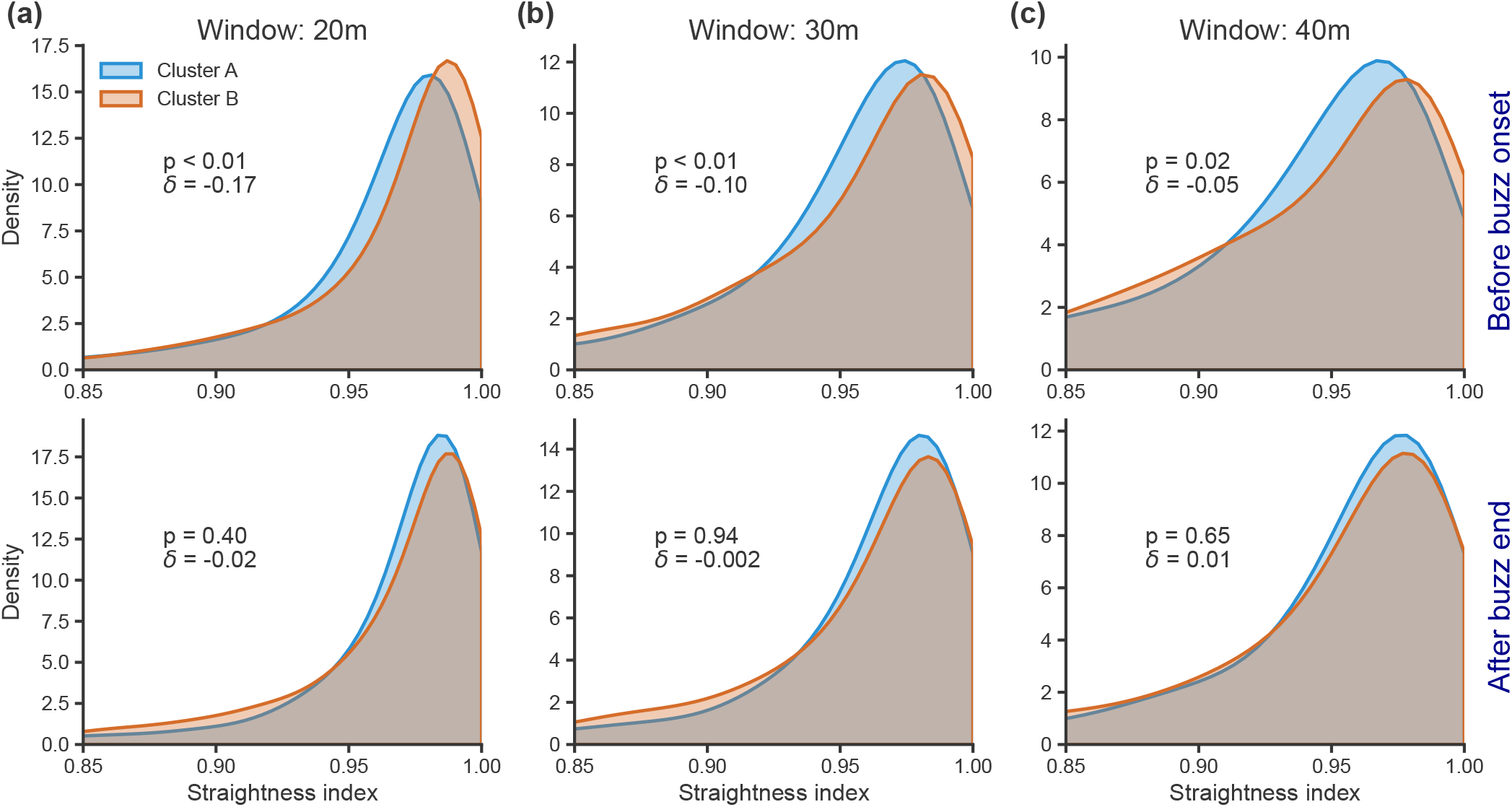
Straightness of movement trajectories before and after prey-capture buzzes. Distributions of the straightness index are shown separately for Type A and Type B buzzes, calculated over path windows immediately preceding buzz onset (top row) and following the last buzz click (bottom row). Window lengths were 20m **(a)**, 30m **(b)** and 40 m **(c)**.

**Figure S18.**
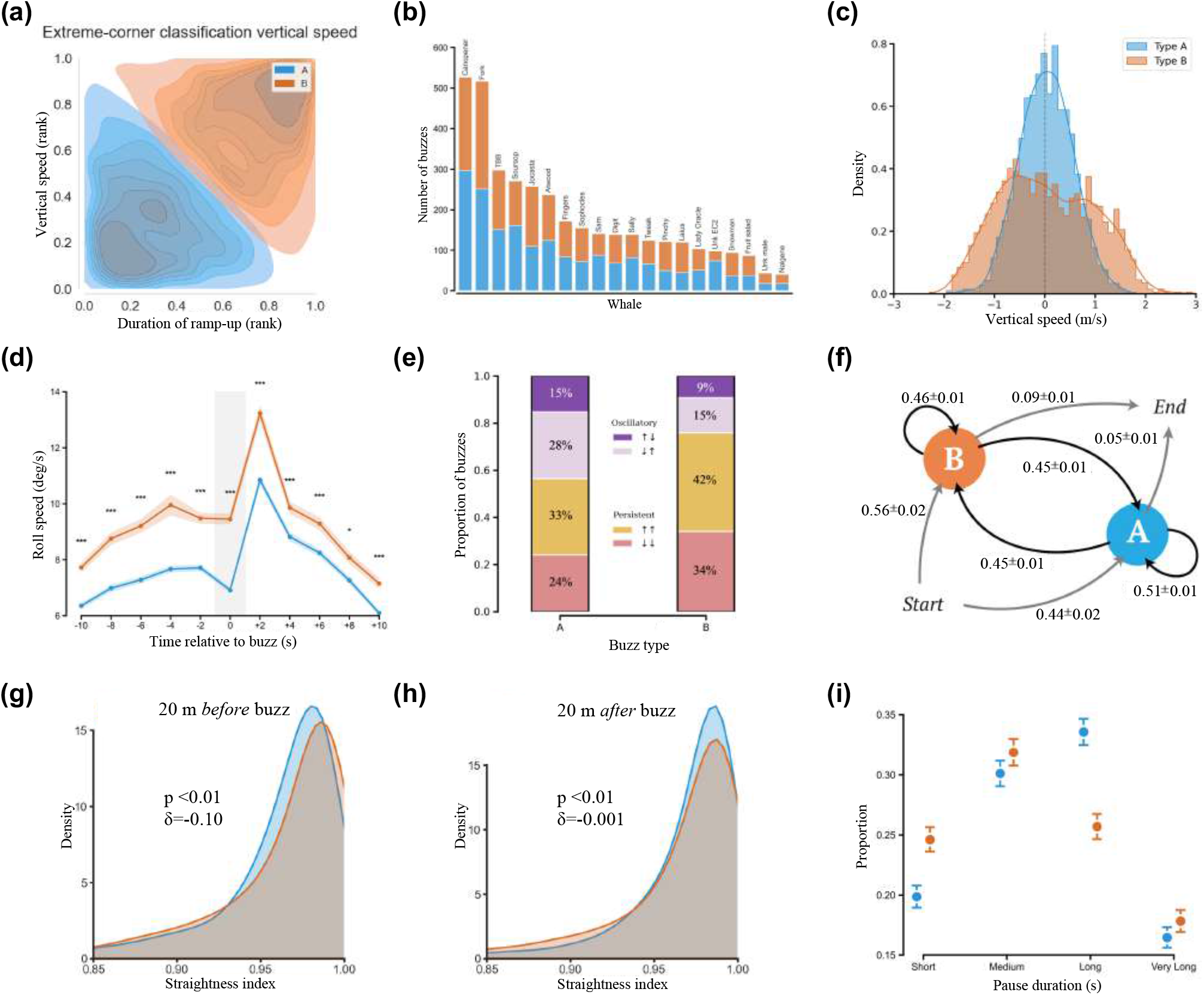
Extreme-corner classification based on vertical speed for the DSWP dataset. **(a)** KDE plots of rank-transformed acoustic and kinematic features (*Duration of ramp-up* vs. *vertical speed*) preserving the bimodal distribution. **(b)** Inter-individual distribution of buzz types, showing that both tactics occur across the sampled whales. **(c)** Distribution of average vertical speeds in the 10 s preceding buzz onset, preserving the unimodal approach for Type A and the bimodal, high-intensity vertical pursuit for Type B. Positive values indicate movement toward the seafloor. **(d)** Mean roll speed (deg/s) calculated in bins of 2 seconds centered on the buzz. Type B events exhibit significantly higher rotational intensity compared to Type A (shaded areas represent standard errors; stars indicate *P <* 0.05, Bonferroni-corrected t-test). **(e)** Proportional distribution of vertical movement classes (oscillatory vs. persistent). Type B attempts show a marked shift toward persistent vertical directionality (76%) compared to the more frequent oscillatory maneuvers in Type A (43%). **(f)** State transition diagram representing buzz sequences as a first-order Markov chain. Arrows indicate transition probabilities (mean *±* s.d. across 200 leave-subset-out resamples) Type B events are approximately twice as likely to terminate a dive (0.09 *±* 0.01) compared to Type A (0.05 *±* 0.01). **(g)** Probability density of the straightness index computed in a 20-meter window preceding buzz onset. Type B pursuits exhibit significantly higher pre-buzz linearity (Mann-Whitney U test, *P <* 0.01; Cliff’s *δ* = −0.10). **(h)** Probability density of the straightness index computed in a 20-meter window following buzz onset. Despite reaching statistical significance (*P <* 0.01), the effect size is negligible (Cliff’s *δ* = −0.001), indicating no meaningful difference between tactics. **(i)** Proportional distribution of pause categories for Type A (blue) and Type B (orange) buzzes. Error bars represent binomial standard errors.

**Figure S19.**
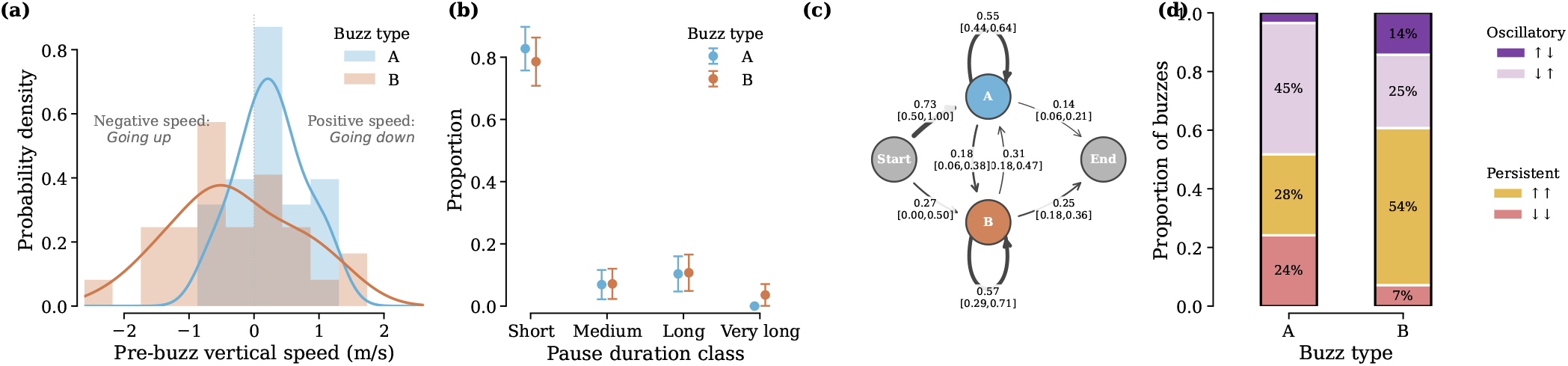
Kinematic and sequential characterization of Type A versus Type B buzzes in the Azorean single-juvenile dataset. Single juvenile sw17_196a; *n* = 58 daytime buzzes classified by the extreme-corner method (*N* = 29 Type A, *N* = 29 Type B). **(a)** Mean signed vertical velocity in the 10 s before buzz onset (positive downwards). Type A was preceded by downward movement (mean +0.25 m s^−1^) and Type B by ascending or near-horizontal movement (mean −0.29 m s^−1^; Mann–Whitney *U* test, *P* = 0.014). **(b)** Proportion of post-buzz pauses in four duration classes; the tactics did not differ (*χ*^2^ = 1.07, *P* = 0.78). Error bars show binomial standard errors. **(c)** First-order Markov chain with 95% bootstrap confidence intervals from 2000 dive resamples. Both tactics showed within-tactic persistence (A→A = 0.55, B→B = 0.57), but the 11 daytime dives preclude precise quantitative comparison with Dominica. **(d)** Vertical-trajectory geometry defined by the signs of vertical velocity before and after each buzz. Type A buzzes were more oscillatory (48%) and Type B more persistent (61%), in the same direction as the DSWP result (44% oscillatory for A and 77% persistent for B; Fig. 3**f**).

## VI. INDEPENDENT AZOREAN VALIDATION

The main-text Azorean analysis uses prey-field echograms to test whether the behavioural extremes identified from ramp-up duration and vertical speed occur under different local prey conditions (Fig. 5). Here we report complementary kinematic and sequence-level measurements from the same 58 daytime buzzes. Because these observations come from a single juvenile and only 11 daytime dives, they are treated as an independent behavioural validation rather than as a population-level estimate or an independent fit of the DSWP mixture model.

The complementary analyses show that the two classes also differ in signed pre-buzz vertical movement and shift in the same direction as DSWP in their broader trajectory geometry: Type A contains more direction reversals, whereas Type B contains more vertically persistent movements (Fig. S19). Both types show within-tactic sequence persistence, although the number of dives is too small for precise comparison of transition probabilities. Post-buzz pause categories do not differ detectably in this dataset. Together with the robust prey-depth difference in the echogram analysis, these results support recurrence of the central acoustic–kinematic contrast while delimiting which secondary sequence features can be evaluated from a single deployment.

## VII. INDEPENDENT VALIDATION IN THE 2024 DOMINICA CETI DEPLOYMENTS

We evaluated whether the core A/B prey-capture contrast was recoverable in independent 2024 Dominica CETI deployments. The available kinematic cohort contained 231 buzzes from 10 tags and 19 dives. Movement data were sampled at 25 Hz and included depth, pitch, roll, and, for five tags, heading. Pitch-adaptive dead reckoning provided three-dimensional speed for all kinematic buzzes; heading-dependent path metrics were restricted to 149 buzzes from the five tags with valid heading. The source structures did not contain prey echograms, and whale identity, age, and sex fields were unavailable, precluding prey-field and per-individual stratified analyses.

The smaller sample did not yield the clear bimodal GMM structure observed in the DSWP dataset, paralleling the limitation encountered in the Azorean analysis. We therefore used the same extreme-corner approach in the rank-transformed ramp-up-duration *×* 3D-speed plane. This classified 115 short-ramp-up, slower buzzes as Type A and 106 long-ramp-up, faster buzzes as Type B (Fig. S20). The purpose of this analysis is not to claim an independent recovery of the mixture model, but to test whether buzzes at the two behavioral extremes carry the same kinematic signatures.

The principal acoustic–kinematic phenotype reproduced. Type B buzzes were faster than Type A by construction of the validation axis and also showed higher mean vertical speed (median 1.16 vs 0.43 m s^−1^; Cliff’s *δ* = −0.78, *P <* 0.001), greater mean roll speed (0.023 vs 0.012 rad s^−1^; *δ* = −0.34, *P <* 0.001), and straighter pre-buzz approaches in the heading-resolved subset (median straightness 0.982 vs 0.975; *δ* = −0.22, *P* = 0.02). Type A buzzes occurred deeper on average (692 vs 645 m; *δ* = 0.25, *P* = 0.002) and were more often associated with oscillatory vertical geometry (48% vs 27%; Fisher’s exact-test odds ratio = 2.43, *P* = 0.002), whereas Type B more often maintained vertical direction (Fig. S21). Thus, the independent deployments recover the same behavioral dichotomy: Type B is fast, high-roll, straight, and vertically persistent, whereas Type A is slower, deeper, and more oscillatory.

These results should be interpreted as an additional validation rather than a population-level replication of every endpoint. The 19 dives provide modest power, and heading-dependent metrics rely on 149 buzzes from five tags. Most importantly, absent individual and demographic metadata prevents testing whether both tactics occur within each identified whale or differ by age and sex. Within these limits, the recovery of the principal kinematic and local path-geometry signatures shows that the A/B contrast is not carried solely by the DSWP clustering method or by the availability of a measured prey field.

**Figure S20.**
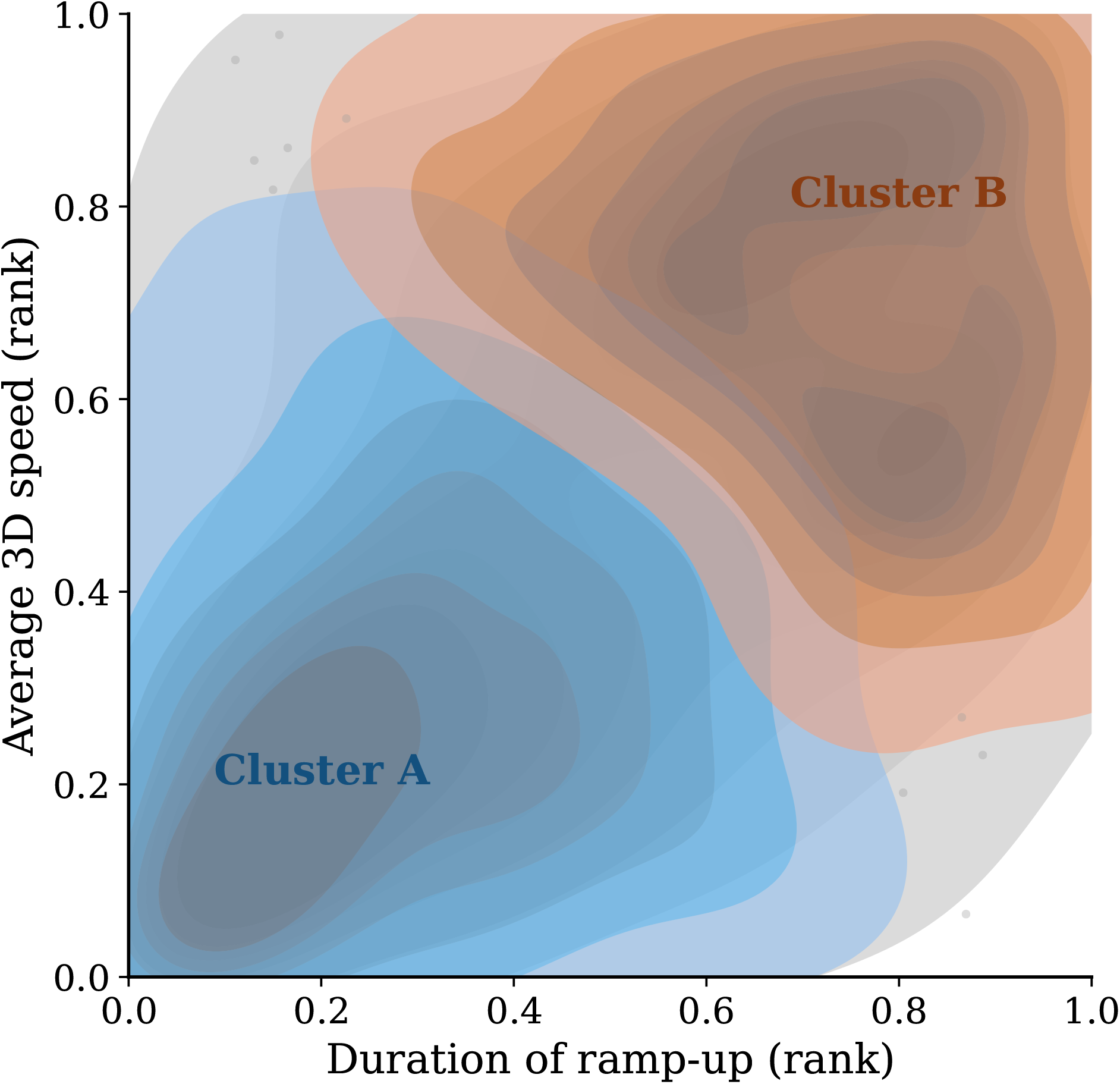
Extreme-corner classification of CETI2024 prey-capture buzzes. Kernel-density contours in the rank-transformed ramp-up-duration *×* pitch-adaptive 3D-speed feature plane. Gray shading shows the full distribution; blue and orange contours show the Type A and Type B extremes used to test the behavioral contrast.

**Figure S21.**
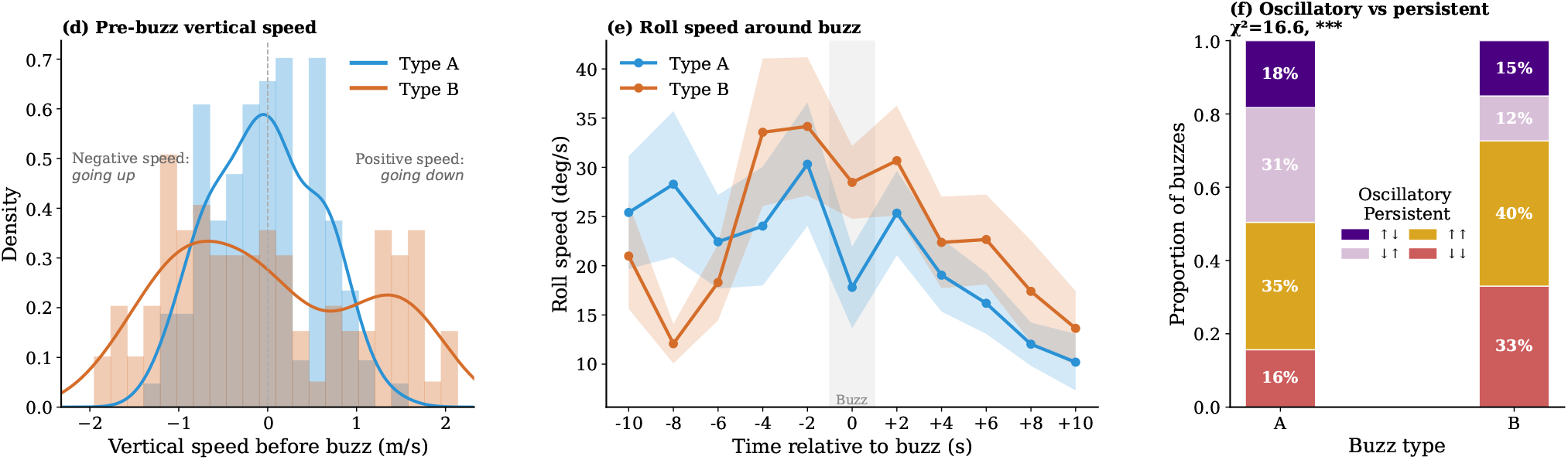
Kinematic validation of the A/B prey-capture contrast in CETI2024. **(a)** Pre-buzz vertical-speed distributions. **(b)** Roll-speed time course around buzz onset, with corrected per-bin comparisons. **(c)** Vertical-trajectory geometry categorized as persistent or oscillatory from the signs of movement before and after each buzz. Across panels, the positive contrast is consistent with Type B representing a faster, higher-roll, more vertically persistent pursuit and Type A a slower, more oscillatory tactic.

## Notes

### Competing Interest Statement

The authors have declared no competing interest.

